# Structural basis for effector transmembrane domain recognition by type VI secretion system chaperones

**DOI:** 10.1101/2020.10.28.356139

**Authors:** Shehryar Ahmad, Kara K. Tsang, Kartik Sachar, Dennis Quentin, Tahmid M. Tashin, Nathan P. Bullen, Stefan Raunser, Andrew G. McArthur, Gerd Prehna, John C. Whitney

## Abstract

Type VI secretion systems facilitate the delivery of antibacterial effector proteins between neighbouring Gram-negative bacteria. A subset of these effectors harbor N-terminal transmembrane domains (TMDs) implicated in effector translocation across the target cell membrane. However, the abundance and distribution of these TMD-containing effectors has remained unknown. Here we report the discovery of prePAAR, a conserved motif found in over 6,000 putative TMD-containing effectors. Based on their differing sizes and number of TMDs these effectors fall into two distinct classes that are unified by their requirement for a member of the Eag family of T6SS chaperones for export. Co-crystal structures of class I and class II effector TMD-chaperone complexes from *Salmonella* Typhimurium and *Pseudomonas aeruginosa*, respectively, reveals that Eag chaperones mimic transmembrane helical packing to stabilize effector TMDs. In addition to participating in the chaperone-TMD interface, we find that prePAAR functions to facilitate proper folding of the downstream PAAR domain, which is required for effector interaction with the T6SS spike. Taken together, our findings define the mechanism of chaperone-assisted secretion of a widespread family of T6SS membrane protein effectors.

## Introduction

Bacteria secrete proteins to facilitate interactions with their surrounding environment. In Gram-negative bacteria, the transport of proteins across cellular membranes often requires the use of specialized secretion apparatuses found within the cell envelope. One such pathway is the type VI secretion system (T6SS), which in many bacterial species functions to deliver antibacterial effector proteins from the cytoplasm directly into an adjacent bacterial cell via a one-step secretion event (Russell et al., 2011). A critical step that precedes type VI secretion is the selective recruitment of effectors to the T6SS apparatus. Recent work has shown that for many effectors this process requires chaperone proteins, which are thought to maintain effectors in a ‘secretion-competent’ state (Unterweger et al., 2017). However, to-date, no molecular-level evidence exists to support this idea.

The T6SS is comprised of two main components: a cell envelope-spanning membrane complex and a cytoplasmic bacteriophage tail-like complex. The latter contains a tube structure formed by many stacked copies of hexameric ring-shaped <u>h</u>emolysin <u>c</u>o-regulated <u>p</u>rotein (Hcp) capped by a single homotrimer of <u>v</u>aline-<u>g</u>lycine <u>r</u>epeat protein <u>G</u> (VgrG)(Mougous et al., 2006, Spinola-Amilibia et al., 2016). Together, these proteins form an assembly that resembles the tail-tube and spike components of contractile bacteriophage (Renault et al., 2018). Additionally, VgrG proteins interact with a single copy of a cone-shaped proline-<u>a</u>lanine-<u>a</u>lanine-<u>a</u>rginine (PAAR) domain-containing protein that forms the tip of the VgrG spike (Shneider et al., 2013). Altogether, PAAR, Hcp and VgrG are necessary for T6SS function, and during a secretion event these components are themselves delivered into target cells (Cianfanelli et al., 2016a). Prior to its export from the cell, the bacteriophage tail-like complex is loaded with toxic effector proteins. In contrast to proteins that are exported by the general secretory pathway, T6SS effectors do not contain linear signal sequences that facilitate their recognition by the T6SS apparatus. Instead, effectors transit the T6SS via physical association with Hcp, VgrG or PAAR proteins (Cianfanelli et al., 2016b).

In addition to its role in effector export, Hcp also possesses chaperone-like properties that facilitate cytoplasmic accumulation of Hcp-interacting effectors prior to their secretion (Silverman et al., 2013). This chaperone activity has been attributed to the interior of the ~4 nm pore formed by hexameric Hcp rings, which are wide enough to accommodate small, single-domain effectors. Individual Hcp rings appear to possess affinity towards multiple unrelated effectors. However, the molecular basis for this promiscuous substrate recognition is unknown.

In contrast to their Hcp-associated counterparts, VgrG-linked effectors are typically comprised of multiple domains and often require effector-specific chaperones for stability and/or to facilitate their interaction with the VgrG spike. Thus far, three effector-specific chaperone families belonging to the DUF1795, DUF2169 and DUF4123 protein families have been described. Studies on representative DUF2169 and DUF4123 proteins indicate that these chaperones minimally form ternary complexes with their cognate effector and a PAAR protein to facilitate the ‘loading’ of the PAAR domain and effector onto their cognate VgrG (Bondage et al., 2016, Burkinshaw et al., 2018). In contrast, DUF1795 proteins, also known as <u>E</u>ffector <u>a</u>ssociated gene (Eag) chaperones, interact with so-called ‘evolved’ PAAR proteins in which the PAAR and toxin domains are found as a single polypeptide chain (Whitney et al., 2015, Alcoforado Diniz and Coulthurst, 2015).

Biochemical characterization of the Eag chaperone EagT6 from *P. aeruginosa* found that this chaperone interacts with TMDs found in the N-terminal loading and translocation region (NLTR) of its associated effector, Tse6 (Quentin et al., 2018). In the presence of lipid vesicles, Tse6 spontaneously inserts into membranes causing EagT6 chaperones to be released suggesting that EagT6 maintains the N-terminal TMDs in a pre-insertion state prior to toxin domain delivery across the inner membrane of target bacteria. However, it is not known whether the ‘solubilization’ of TMDs in aqueous environments represents a general role for Eag chaperones and if so, it is unclear how they maintain effector TMDs in a pre-insertion state.

In this work, we report the identification of prePAAR, a highly conserved motif that enabled the identification of over 6,000 putative T6SS effectors, all of which possess N-terminal TMDs and co-occur in genomes with Eag chaperones. Further informatics analyses found that these candidate effectors can be categorized into one of two broadly defined classes. Class I effectors belong to the Rhs family of proteins, are comprised of ~1200 amino acids and possess a single region of N-terminal TMDs. Class II effectors are ~450 amino acids in length and possess two regions of N-terminal TMDs. We validate our informatics approach by showing that a representative member of each effector class requires a cognate Eag chaperone for T6SS-dependent delivery into susceptible bacteria. Crystal structures of Eag chaperones in complex with the TMDs of cognate class I and class II effectors reveal the conformation of effector TMDs prior to their secretion and insertion into target cell membranes. In addition to participating in chaperone-effector interactions, structure-guided mutagenesis of hydrophilic residues within prePAAR show that this motif also catalyzes the appropriate folding of the downstream PAAR domain, enabling its interaction with its cognate VgrG. Collectively, our data provide the first high-resolution structural snapshots of T6SS effector-chaperone interactions and define the molecular determinants for effector TMD stabilization and recruitment to the T6SS apparatus.

### prePAAR is a motif found in TMD-containing effectors that interact with Eag chaperones

Characterization of Eag chaperones and their associated effectors has thus far been limited to the EagT6-Tse6 and EagR1-RhsA chaperone-effector pairs from *P. aeruginosa* and *Serratia marcescens*, respectively (Cianfanelli et al., 2016a, Whitney et al., 2015). In both cases, the chaperone gene is found upstream of genes encoding its cognate effector and an immunity protein that protects the toxin-producing bacterium from self-intoxication (Figure 1A). We previously showed that EagT6 interacts with the N-terminal TMDs of Tse6, an observation that led us to hypothesize a general role for Eag chaperones in ‘solubilizing’ hydrophobic TMDs of effectors in the aqueous environment of the cytoplasm so they can be loaded into the T6SS apparatus (Figure 1B) (Quentin et al., 2018). However, evidence supporting this general role is lacking because homology-based searches for additional Eag chaperones can yield difficult to interpret results due to a scarcity of conserved residues and homology of this protein family to the phage protein DcrB (Samsonov et al., 2002), which is widely distributed in both T6SS-positive and T6SS-negative organisms. Similarly, the identification of N-terminal TMD-containing PAAR effectors that might require Eag chaperones is also challenging because many PAAR domain-containing effectors lack TMDs (Shneider et al., 2013), and aside from being comprised of hydrophobic residues, the TMDs themselves are poorly conserved.

**Figure 1.**
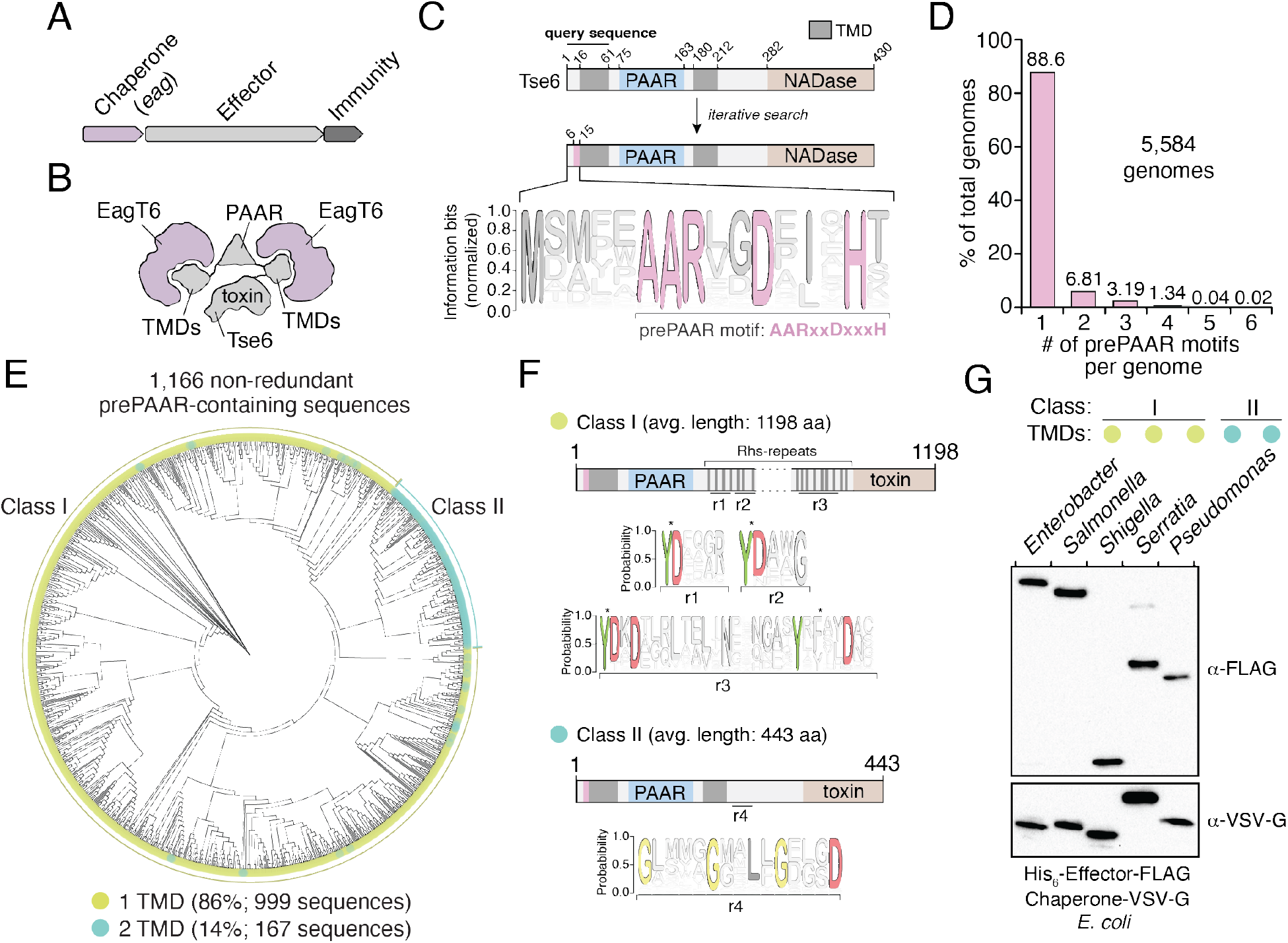
The prePAAR motif is conserved across multiple bacterial genera and is found in T6SS effectors that interact with Eag chaperones. A) Genomic arrangement of T6SS chaperone-effector-immunity genes for characterized *<u>e</u>ffector <u>a</u>ssociated <u>g</u>ene* family members (*eag*; shown in blue), which encode DUF1795 domain-containing chaperones. B) Schematic depicting Eag chaperone interactions with the transmembrane domain (TMD) regions of the model chaperone-effector pair EagT6-Tse6. C) Protein architecture and sequence logo for the prePAAR motif found in the N-terminus of Tse6. An alignment of 2,054 sequences was generated using the 61 N-terminal residues of Tse6 as the search query. The relative frequency of each residue and information content in bits was calculated at every position of the sequence and then normalized by the sum of each position’s information bits. Transparency is used to indicate probability of a residue appearing at a specific position. Residues coloured in pink correspond to the prePAAR motif: AARxxDxxxH. D) Genomes from genera of Proteobacteria known to contain functional T6SSs (*Burkholderia*, *Escherichia*, *Enterobacter*, *Pseudomonas*, *Salmonella*, *Serratia*, *Shigella*, *Yersinia*) were screened for unique prePAAR effectors. Percentage of total genomes that contained 1 to 6 prePAAR motifs is indicated. E) Phylogenetic distribution of 1,166 non-redundant prePAAR-containing effectors identified in **B**. TM prediction algorithms were used to quantify the number of TM regions in each effector. The two classes that emerged are labeled in green (class I; 1 TM region-containing effectors) and blue (class II; 2 TM region-containing effectors). Branch lengths indicates evolutionary distances. F) Effector sequences within class I or class II were aligned, and a sequence logo was generated based on the relative frequency of each residue at each position to identify characteristic motifs of both classes. Four different regions (r1-r4) after the PAAR and TM regions were found to harbour conserved residues. Class I effectors contain YD repeat regions (r1-3) characteristic of Rhs proteins whereas a GxxxxGxxLxGxxxD motif (r4) was identified in class II effectors. G) Western blot analysis of five effector-chaperone pairs that were selected from the indicated genera, based on the analysis in **B**. Each pair was co-expressed in *E. coli* and co-purified using nickel affinity chromatography. The class and number of TM regions from each pair are indicated. Locus tags for each pair (e, effector; c, chaperone) are as follows: *Enterobacter* (e: ECL_01567, c: ECL_01566), *Shigella* (e: SF0266, c: SF3490), *Salmonella* (e: SL1344_0286, c: SL1344_0285), *Serratia* (e: Spro_3017, c: Spro_3016), *Pseudomonas* (e: PA0093, c: PA0094) . Note that the Rhs component of the class I prePAAR effector SF0266 is encoded by the downstream open reading frame SF0267 (see Extended Data Figure 1C for details).

In an attempt to overcome the challenges associated with identifying Eag-interacting T6SS effectors, we used *jackhmmer* to generate a sequence alignment hidden Markov model (HMM) for the N-terminal 60 residues of Tse6 using an iterative search procedure that queried the UniProtKB database (Johnson et al., 2010). We reasoned that if there exists a molecular signature present in effector proteins indicative of Eag chaperone association, it would be located within this region of Tse6 homologous proteins because it contains a known chaperone binding site. Remarkably, the HMM we obtained revealed a nearly invariant AARxxDxxxH motif, which in Tse6 is found in the first 15 residues of the protein and is immediately N-terminal to its first TMD (Figure 1C). In total, our query identified over 2,054 proteins containing this motif (Table 1 and Figure 1—figure supplement 1A). Among these candidate effectors, our search identified the recently characterized toxins Tre1, Tas1, DddA as well as many toxins of unknown function indicating that our approach may have identified T6SS effectors with novel biochemical activities (Ting et al., 2018, Ahmad et al., 2019, Mok et al., 2020). Interestingly, prior to any knowledge of PAAR domains or Eag chaperones being involved in T6SS function, Zhang and colleagues noted the existence of this N-terminal motif in an informatics analysis of bacterial nucleic acid degrading toxins (Zhang et al., 2011). Here, they refer to it as prePAAR because PAAR sequences were found C-terminal to the motif. We have adhered to this name because as described in detail below, this pattern holds true for the thousands of candidate effectors identified in our search.

Examination of our putative effector sequences revealed that prePAAR is substantially enriched in bacterial genera with characterized T6SSs including *Pseudomonas*, *Burkholderia*, *Salmonella*, *Shigella*, *Escherichia*, *Enterobacter*, *Yersinia*, and *Serratia*. Interestingly, no prePAAR motifs were identified in *Vibrio* despite an abundance of species within this genus possessing highly active bacteria-targeting T6SSs. We next obtained all 56,324 available genomes from NCBI for the abovementioned genera and found that 26,327 genomes encode at least one prePAAR motif. After removing all redundant sequences, 6,129 unique prePAAR-containing proteins present across 5,584 genomes were used for further analyses (Table 2, ‘unfiltered’ dataset). In these genomes, we determined that approximately 90% encode a single prePAAR motif, although instances where prePAAR is present up to six times within a single genome were also identified (Figure 1D). To determine if these unique proteins are probable TMD-containing T6SS effectors that require Eag chaperones for secretion, we next examined each prePAAR-containing protein and its associated genome for the following three criteria: 1) the existence of an Eag chaperone encoded in the same genome, 2) the presence of a downstream PAAR domain and 3) predicted TMDs in the first 300 amino acids of the protein (Krogh et al., 2001, Kall et al., 2007). The location restriction in our TMD search was used in order to exclude C-terminal toxin domains that possess TMDs, which differ from N-terminal translocation TMDs in that they may not require chaperones for secretion (Mariano et al., 2019). We searched each genome for Eag proteins using an HMM for DUF1795 and found that 99.5% (5,554/5,584) of prePAAR-containing genomes also possessed at least one *eag* gene (Jones et al., 2014). In approximately 14% of the 5,554 genomes analyzed, the number of prePAAR motifs matched the number of Eag homologues. In the remainder of cases, the number of Eag homologous proteins exceeded the number of prePAAR motifs, with a weighted average of 2.5 paralogues per genome. As is the case with *eagT6-tse6* and *eagR1-rhsA*, ~90% of the identified prePAAR-containing effector genes appear directly beside an *eag* gene whereas the remaining ~10% are found in isolation suggesting that their putative chaperone is encoded elsewhere in the genome. We removed pre-PAAR-containing protein fragments (proteins less than 100 amino acids in length) and further reduced redundancy by clustering sequences with 95% identity. Remarkably, in all but two of the remaining 1,166 prePAAR-containing proteins, we identified a PAAR domain, indicating a probable functional relationship between prePAAR and PAAR. The two prePAAR-containing proteins lacking a PAAR domain were either adjacent to a gene encoding a PAAR domain-containing protein or directly beside T6SS structural genes. Finally, we searched 1,166 prePAAR-containing proteins for TMDs and found that all protein sequences contained predicted TMDs with 86% having one region of TMDs and 14% having two regions of TMDs. In sum, our prePAAR-based search procedure identified thousands of candidate effector proteins possessing properties consistent with the requirement for an Eag chaperone for T6SS-dependent export.

To further analyze our collection of prePAAR-containing effectors, we built a phylogenetic tree from 1,166 non-redundant effector sequences that represent the diversity present in our collection of sequences (Fig. 1E). Interestingly, two distinct sizes of proteins emerged from this analysis: large prePAAR effectors that are over 1000 amino acids in length and small prePAAR effectors comprised of 350-450 amino acids (Figure 1E and Figure 1—figure supplement 1B). As noted previously, all effectors contained predicted TMDs; however, large effectors almost exclusively contained a single region of TMDs N-terminal to their PAAR domain whereas most small effectors contained TMD regions N- and C-terminal to their PAAR domain. To distinguish between these two domain architectures, we hereafter refer to large, single TMD region-containing prePAAR effectors as class I and small, two TMD region-containing prePAAR effectors as class II. Notably, class I effectors also contain numerous YD repeat sequences, which are a hallmark of rearrangement <u>h</u>ot<u>s</u>pot (Rhs) proteins that function to encapsulate secreted toxins (Figure 1F)(Busby et al., 2013). Conversely, class II effectors are distinguished by a GxxxxGxxLxGxxxD motif in addition to their second TMD region.

As a first step towards validating our informatics approach for identifying Eag chaperone-effector pairs, we assessed the ability of several newly identified Eag chaperones to interact with the prePAAR-containing effector encoded in the same genome. We previously demonstrated that the class II effector Tse6 interacts with EagT6 and we similarly found that when expressed in *E. coli*, Eag chaperones from *Enterobacter cloacae*, *Salmonella* Typhimurium, *Shigella flexneri* and *Serratia proteamaculans* co-purified with their predicted cognate effector (Figure 1G and Figure 1—figure supplement 1C). Collectively, these findings indicate that prePAAR proteins constitute two classes of TMD-containing T6SS effectors and that representative members from both classes interact with Eag chaperones.

### Eag chaperones are specific for cognate prePAAR effectors

We next sought to examine the specificity of Eag chaperones towards prePAAR effectors in a biologically relevant context. To accomplish this, we inspected our list of prePAAR effectors and found that the soil bacterium *Pseudomonas protegens* Pf-5 possesses both a class I and class II effector, encoded by the previously described effector genes *rhsA* and *tne2*, respectively (Tang et al., 2018). Furthermore, the genome of this bacterium encodes two putative Eag chaperones, PFL_6095 and PFL_6099, which have 25% sequence identity between them (Figure 2A). PFL_6095 is found upstream of *rhsA* and is likely co-transcribed with this effector whereas PFL_6099 is not found next to either effector gene. To examine the relationship between these genes, we generated strains bearing single deletions in each effector and chaperone gene and conducted intraspecific growth competition assays against *P. protegens* recipient strains lacking the *rhsA-rhsI* or *tne2-tni2* effector-immunity pairs. We noted that protein secretion by the T6SS of *P. protegens* is substantially inhibited by the threonine phosphorylation pathway, so we additionally inactivated the threonine phosphatase encoding gene *pppA* in recipients to induce a ‘tit-for-tat’ counterattack by wild-type donor cells (Figure 2—figure supplement 1A-B)(Mougous et al., 2007, Basler et al., 2013). Consistent with the effector-immunity paradigm for bacteria-targeting T6SSs, wild-type *P. protegens* readily outcompeted Δ*rhsA* Δ*rhsI* Δ*pppA* and Δ*tne2* Δ*tni2* Δ*pppA* strains in a *rhsA*- and *tne2*-dependent manner, respectively (Figure 2B). Additionally, we found that a strain lacking PFL_6095 no longer exhibited a co-culture fitness advantage versus a Δ*rhsA* Δ*rhsI* Δ*pppA* recipient but could still outcompete *tne2* sensitive recipients to the same extent as the wild-type strain. Conversely, a ΔPFL_6099 strain outcompeted Δ*rhsA* Δ*rhsI* Δ*pppA* but not Δ*tne2* Δ*tni2* Δ*pppA* recipients. Together, these data indicate that the delivery of RhsA and Tne2 into susceptible target cells requires effector-specific *eag* genes.

**Figure 2.**
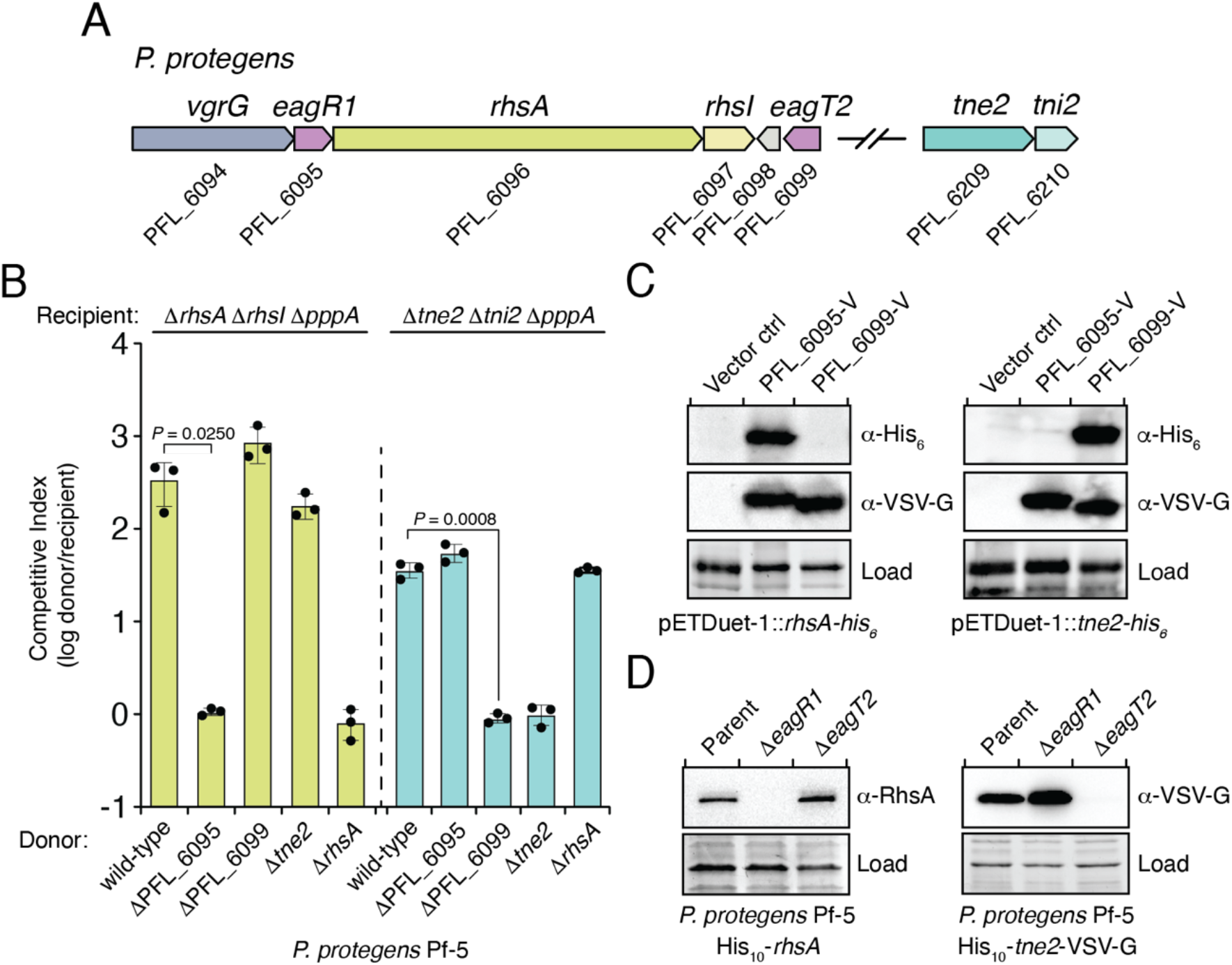
Eag chaperones are specific for their cognate prePAAR effector and are necessary for effector stability *in vivo*. A) Genomic context of two prePAAR-containing effector-immunity pairs from *P. protegens* Pf-5. RhsA is a class I effector (shown in green) and Tne2 is a class II effector (shown in blue). Shading is used to differentiate effector (dark) and immunity genes (light). Predicted *eag* genes are shown in purple. B) Outcome of intraspecific growth competition assays between the indicated *P. protegens* donor and recipient strains. Donor strains were competed with recipient strains lacking *rhsA-rhsI* (green) or *tne2-tni2* (blue). Both recipients are lacking *pppA* to stimulate type VI secretion. Data are mean ±s.d. for *n* = 3 biological replicates and are representative of two independent experiments; *P* values shown are from two-tailed, unpaired *t*-tests. C) Western blot analysis of *E. coli* cell lysates from cells expressing the indicated effectors (RhsA or Tne2) and either empty vector, PFL_6095-V or PFL_6099-V. D) Affinity-tagged RhsA or Tne2 were purified from cell fractions of the indicated *P. protegens* strains and visualized using western blot analysis. Deletion constructs for each *eag* gene were introduced into each of the indicated parent backgrounds. A non-specific band present in the SDS-PAGE gel was used as a loading control. C-D) Data are representative of two independent experiments.

To test the ability of PFL_6095 and PFL_6099 to act as RhsA- and Tne2-specific chaperones, respectively, we co-expressed each chaperone-effector pair in *E. coli* and examined intracellular effector levels by western blot. Consistent with functioning to promote cognate effector stability, accumulation of RhsA only occurred in the presence of PFL_6095 whereas Tne2 accumulated in cells containing PFL_6099 (Figure 2C). We next examined the stability-enhancing properties of PFL_6095 and PFL_6099 when expressed at native levels in *P. protegens*. Due to challenges associated with detecting RhsA and Tne2 in unconcentrated cell lysates, we constructed chromosomally encoded N-terminal decahistidine-tagged (his_10_) fusions of RhsA and Tne2 to facilitate the enrichment of these proteins from *P. protegens* and confirmed that these fusions did not compromise the ability of these effectors to intoxicate recipients (Figure 3—figure supplement 1A-B). Following affinity purification, RhsA and Tne2 levels were assessed using RhsA and vesicular stomatitis virus glycoprotein epitope (VSV-G) antibodies, respectively. In line with our data in *E. coli*, we were unable to detect RhsA in the absence of PFL_6095 whereas Tne2 was absent in a strain lacking PFL_6099 (Figure 2D). Collectively, these data suggest that Eag chaperones exhibit a high degree of specificity for their cognate effectors. Based on our characterization of these genes, we propose to rename PFL_6095 and PFL_6099 to *eagR1* and *eagT2*, respectively, to reflect their newfound role as chaperones for the prePAAR-containing effectors RhsA and Tne2.

**Figure 3.**
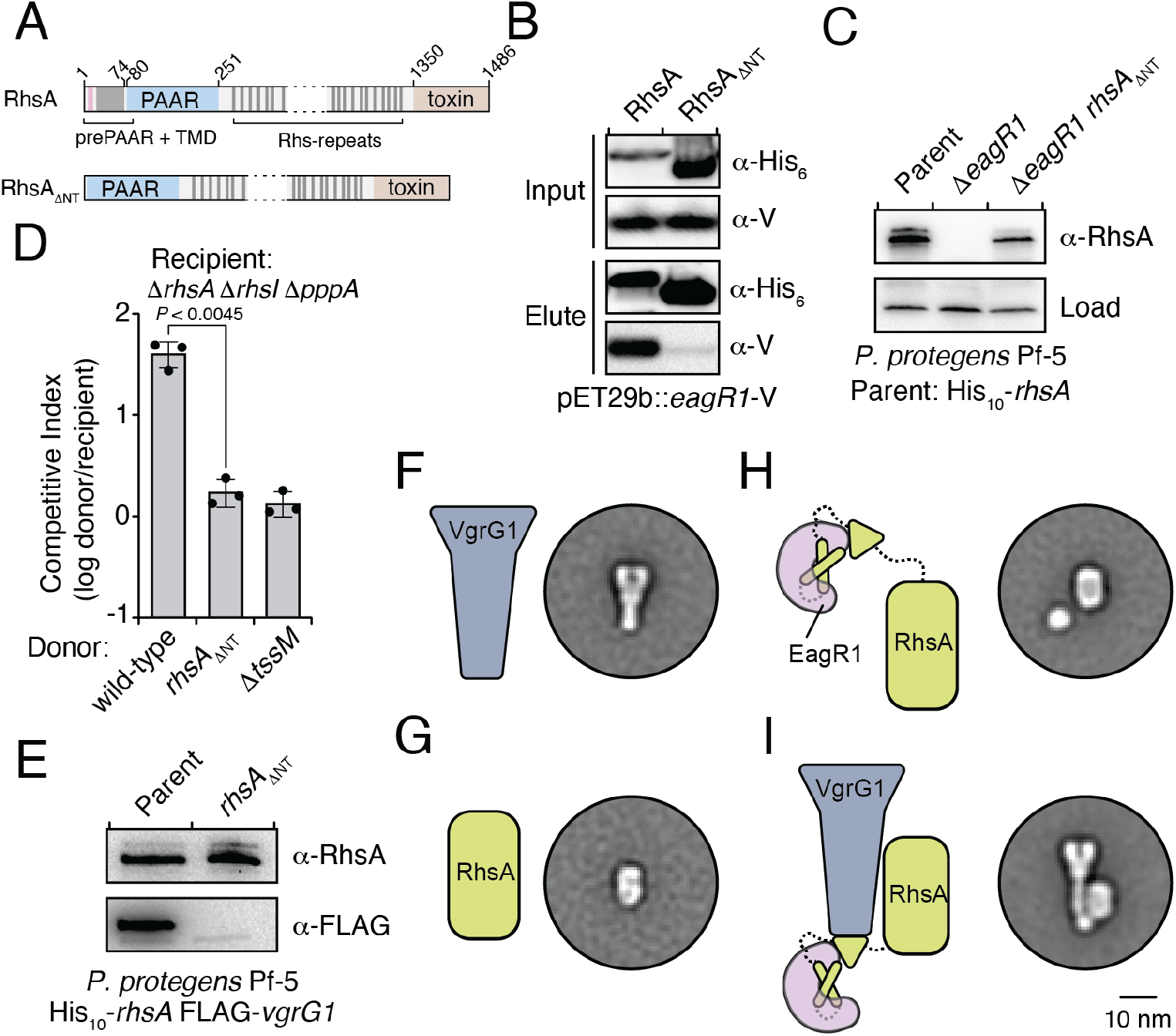
An Eag chaperone promotes the stability of its cognate class I prePAAR effector by interacting with its prePAAR and TMD-containing N-terminus. A) Domain architecture of *P. protegens* RhsA and a truncated variant lacking its prePAAR and TMD-containing N-terminus (RhsA_ΔNT_). B) EagR1 interacts with the N-terminus of RhsA. His_6_-tagged RhsA or RhsA_ΔNT_ and co-expressed with EagR1 in *E. coli*, purified using affinity chromatography and detected by western blot. C) Affinity purification of chromosomally His_10_-tagged RhsA or RhsA_ΔNT_ from cell fractions of the indicated *P. protegens* strains. The parent strain expresses chromosomally encoded His_10_-tagged RhsA. The loading control is a non-specific band on the blot. D) Outcome of growth competition assays between the indicated donor and recipient strains of *P. protegens*. Data are mean ±s.d. for *n* = 3 biological replicates; *P* value shown is from a two-tailed, unpaired *t*-test. E) Affinity purification of His_10_-RhsA or His_10_-RhsA_ΔNT_ from a *P. protegens* Pf-5 strain containing a chromosomally encoded FLAG epitope tag fused to *vgrG1*. FLAG-tagged VgrG1 was detected by western blot. F-I) Representative negative-stain EM class averages for purified VgrG1 (F), RhsA_ΔNT_ (G), EagR1-RhsA complex (H) and EagR1-RhsA-VgrG complex (I). Scale bar represents 10 nm for all images. All proteins were expressed and purified from *E. coli*. B-C, E) Data are representative of two independent experiments.

Previous biochemical studies on the class II prePAAR effector Tse6 and its cognate chaperone EagT6 demonstrated that the two TMD regions of this effector each require an EagT6 chaperone for stability (Quentin et al., 2018). These findings suggest that there may exist a physical limitation to the number of TMDs that a single EagT6 chaperone can stabilize. Our finding that class I prePAAR effectors contain only one TMD region suggests that these proteins may only possess one Eag interaction site (Figure 3A). To test this, we constructed a RhsA variant lacking its N-terminal region (RhsA_ΔNT_) and co-expressed this truncated protein with EagR1 in *E. coli*. Consistent with our hypothesis, affinity purification of RhsA_ΔNT_ showed that this truncated variant does not co-purify with EagR1 (Figure 3B). Additionally, expression of the 74 residue N-terminal fragment of RhsA in isolation was sufficient for EagR1 binding (Figure 3—figure supplement 1C). Our data also demonstrate that in contrast to wild-type RhsA, RhsA_ΔNT_ is stable in the absence of EagR1 when expressed in *E. coli* indicating that the N-terminus imparts instability on the protein in the absence of its cognate chaperone. In *P. protegens*, we could readily detect *rhsA*ΔNT in a strain lacking *eagR1*, corroborating our findings in *E. coli* (Figure 3C). However, despite the enhanced stability of chaperone ‘blind’ RhsA_ΔNT_, a *P. protegens* strain expressing this truncation was no longer able to outcompete RhsA-sensitive recipient cells demonstrating an essential role for the chaperone-bound N-terminus during interbacterial competition (Figure 3D).

After ruling out the possibility that truncating the N-terminus of RhsA affects the growth-inhibitory activity of its C-terminal toxin domain (Figure 3—figure supplement 1D), we next examined the ability of RhsA_ΔNT_ to interact with its cognate secreted structural component of the T6SS apparatus. T6SS effectors encoded downstream of *vgrG* genes typically rely on the encoded VgrG protein for delivery into target cells (Whitney et al., 2014). Consistent with this pattern, PFL_6094 encodes a predicted VgrG protein, herein named VgrG1, which we confirmed is required for RhsA-mediated growth inhibition of susceptible target cells (Figure 3—figure supplement 1E). Furthermore, using a *P. protegens* strain expressing His_10_-tagged RhsA and FLAG-tagged VgrG1 from their native loci, we found that these proteins physically interact to form a complex (Figure 3E). To test if the absence of the chaperone-bound N-terminus affects the formation of this complex, we used our *E. coli* co-expression system to purify RhsA-EagR1-VgrG1 complexes. These experiments show that RhsA_ΔNT_ is not able to interact with VgrG1, even though this truncated protein possesses its PAAR domain, which in T6SS effectors lacking prePAAR and TMDs in their N-terminus, is sufficient for VgrG interaction (Figure 3— figure supplement 1F)(Bondage et al., 2016).

To gain insight into how EagR1 binding facilitates RhsA interaction with VgrG1, we next performed negative-stain electron microscopy (EM) to examine the configuration of each subunit within this complex. To facilitate the accurate identification of each component, we obtained class averages of purified VgrG1, RhsA_ΔNT_, RhsA-EagR1 complex and RhsA-EagR1-VgrG1 complex (Figure 4—figure supplement 1A-H). As expected, isolated VgrG1 and RhsA_ΔNT_ proteins appeared as characteristic spike- and barrel-shaped proteins, respectively (Spinola-Amilibia et al., 2016, Busby et al., 2013); Figures 3F and 3G). Intriguingly, images of RhsA-EagR1 complexes contained a sphere-shaped object that likely represents a subcomplex between EagR1 and the N-terminus of RhsA (Figure 3H). Lastly, the class-averages of RhsA-EagR1-VgrG1 complexes revealed a close association of EagR1 and RhsA with the tip of the VgrG spike, which is likely mediated by the PAAR domain of RhsA (Figure 3I). Interestingly, though both complexes exhibit significant rotational flexibility, the average distance between the subcomplex formed by EagR1 and the N-terminus of RhsA is substantially greater in the absence of VgrG1 (average distance: 2.68 nm, *n* = 27 classes versus 1.20 nm, *n* = 26 classes) (Figure 4—figure supplement 1F-H). When taken together with our biochemical experiments, these structural data indicate that EagR1 stabilizes the N-terminus of RhsA, which may also orient the effector such that it can interact with its cognate VgrG.

**Figure 4.**
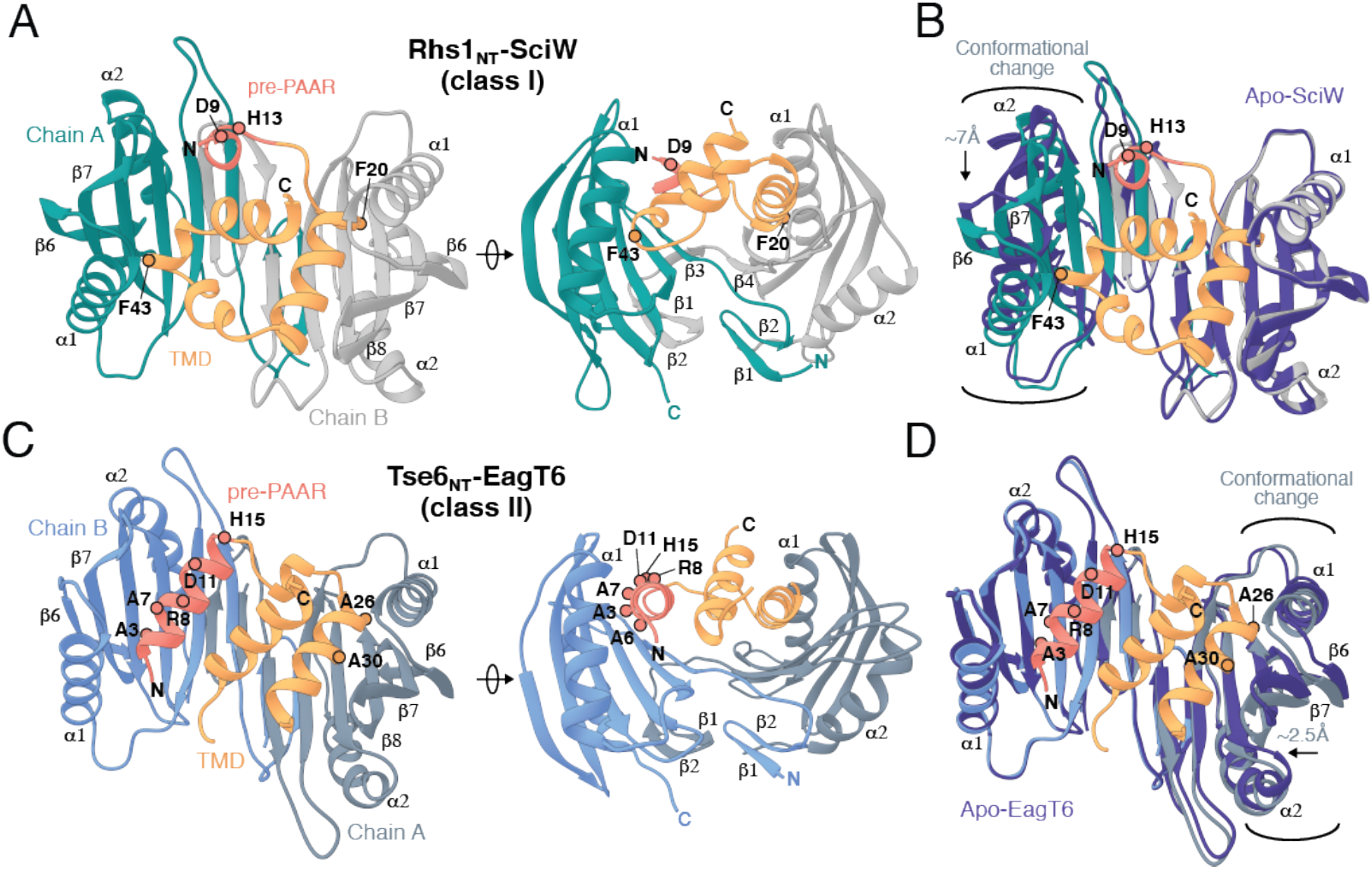
Co-crystal structures of the N-terminus of class I and class II prePAAR effectors in complex with their cognate Eag chaperones. A) An X-ray crystal structure of the Eag chaperone SciW bound to the N-terminus of *Salmonella* Typhimurium class I prePAAR effector Rhs1 (Rhs1_NT_, residues 8-57 are modeled) shown in two views related by a ~90° rotation. B) Structural overlay of the apo-SciW structure with SciW-Rhs1_NT_ complex demonstrates that a considerable conformational change in SciW occurs upon effector binding. C) An X-ray crystal structure of the Eag chaperone EagT6 bound to the N-terminus of Tse6 (Tse6_NT_, residues 1-38 and 41-58 are modeled) shown in two views related by a ~90° rotation. D) Structural overlay of the apo-EagT6 structure (PDB 1TU1) with the EagT6-Tse6_NT_ complex shows a minor conformational change in EagT6 occurring upon effector binding. Eag chaperones are colored by chain, N-terminal transmembrane domains (TMDs) are colored in orange, the pre-PAAR motif in red, and apo chaperone structure in dark blue. Positions of residues of interest in the effector N-terminal regions are labeled.

### Eag chaperones bind effector TMDs by mimicking transmembrane helical packing

In addition to a TMD-containing region, the N-terminus of prePAAR effectors also harbours the prePAAR motif itself. However, the negative stain EM images of RhsA-EagR1-VgrG1 particles presented herein and our previously determined single-particle cryo-EM structure of a complex containing Tse6-EagT6-VgrG1 are of insufficient resolution to resolve the structures of chaperone-bound effector TMDs or the prePAAR motif (Quentin et al., 2018). Therefore, to better understand the molecular basis for chaperone-TMD interactions and to gain insight into prePAAR function we initiated X-ray crystallographic studies on both class I and class II effector-chaperone complexes. Efforts to co-crystallize *P. protegens* EagR1 with the prePAAR and TMD-containing N-terminus of RhsA were unsuccessful. However, the EagR1 homologue SciW from *Salmonella* Typhimurium crystallized in isolation and in the presence of the N-terminus of the class I prePAAR effector Rhs1 (Rhs1_NT_), allowing us to determine apo and effector bound structures to resolutions of 1.7Å and 1.9Å, respectively (Figure 4 and Table 4). Similar to RhsA, we confirmed that a Rhs1_ΔNT_ variant was unable to bind its cognate chaperone, SciW (Figure 3—figure supplement 1G). The structure of the EagT6 chaperone was previously solved as part of a structural genomics effort and we were additionally able to obtain a 2.6Å co-crystal structure of this chaperone in complex with the N-terminal prePAAR and first TMD region of the class II effector Tse6 (Tse6_NT_) (Figure 4 and Table 4).

**Table 4.**
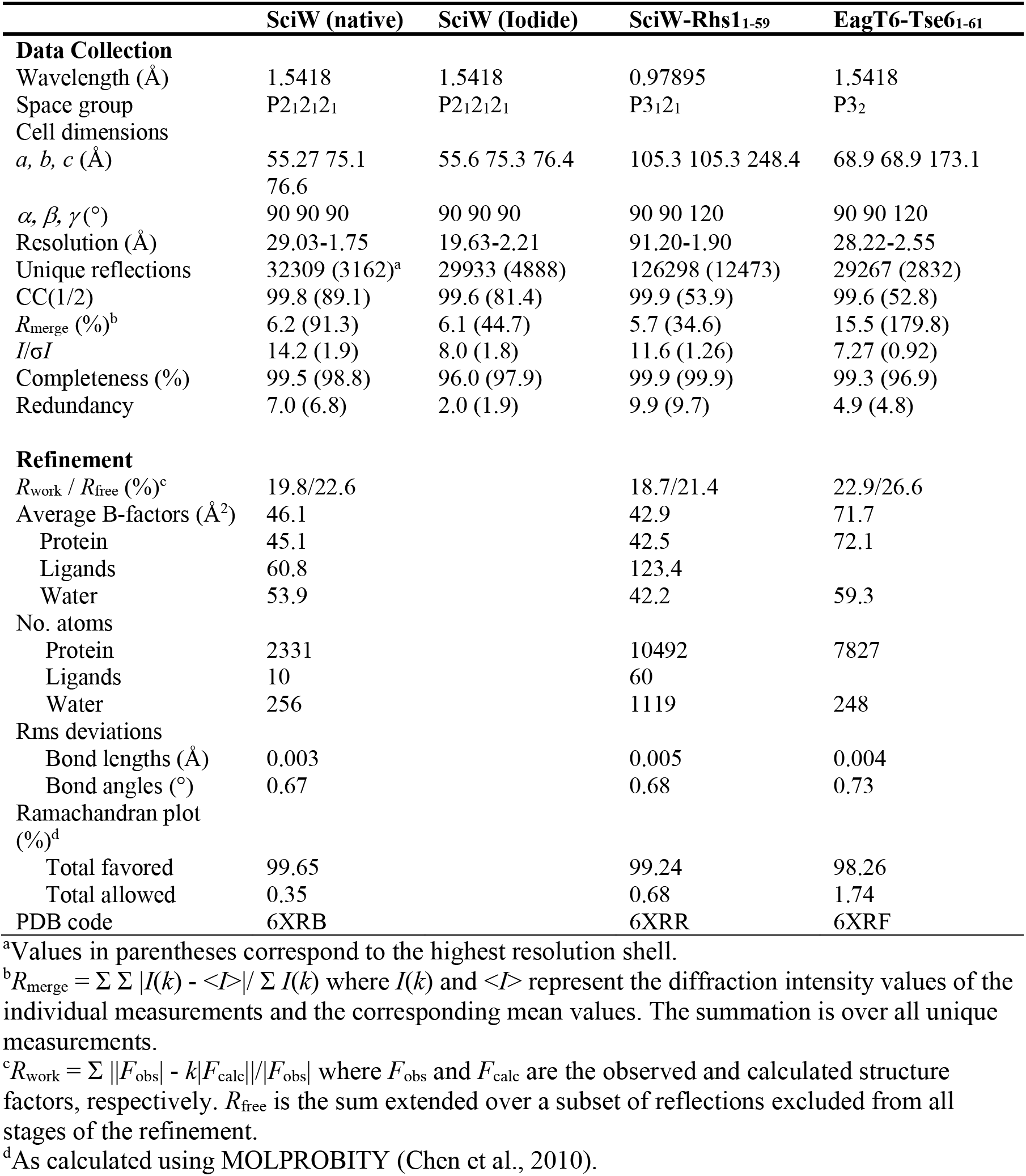
X-ray data collection and refinement statistics.

The overall structure of SciW reveals a domain-swapped dimeric architecture that is similar to the previously described apo structure of *P. aeruginosa* EagT6 though each chaperone differs in its electrostatic surface properties (Figure 5—figure supplement 1A-D) (Whitney et al., 2015). A comparison of the chaperone structures in their apo and effector bound states shows that upon effector binding, both chaperones ‘grip’ the prePAAR-TMD region of their cognate effector in a claw-like manner (Figure 4A-D). Although our biochemical data indicate that Eag chaperones exhibit a high degree of specificity for their associated effector, the internal surface of the claw-shaped dimer contains a number of conserved residues that make critical interactions with the TM helices in both complexes (Figure 5A-F). For example, I22 and I24 of EagT6 create a hydrophobic surface in the ‘palm’ of the claw, which is flanked on either side by symmetrical hydrophobic surfaces comprised of A62, L66, L98, F104 and I113 (Figure 5B-D). Furthermore, the conserved hydrophilic residues S37, S41, Q58, and Q102 also interact with the bound effectors by making bifurcated hydrogen bonds to amide or carbonyl groups in the peptide backbone of the TM helices (Figure 5E and 5F). These polar interactions between chaperone and effector TM helices are striking because they are reminiscent of polar interactions seen within the helical packing of alpha helical transmembrane proteins, which often use serine and glutamine residues to mediate inter-helical interactions via bifurcated hydrogen bonds between side-chain and main-chain atoms (Dawson et al., 2002, Dawson et al., 2003, Adamian and Liang, 2002). Additionally, EagT6 and SciW provide ‘knob-hole-like’ interactions, which also feature prominently in membrane protein packing (Curran and Engelman, 2003). Knob-hole interactions involve a large hydrophobic residue on one TM helix acting as a ‘knob’ to fill the hole provide by a small residue such as glycine or alanine on another TM-helix. TM holes are typically created by GxxxG/A motifs such as those found in G19-A24 (Rhs1) and G25-A30 (Tse6). In this case, the conserved Eag chaperone residue L66 provides a knob for the A24/30 hole (Figure 5E and 5F). Given that the Eag chaperone dimer creates a hydrophobic environment with complementary knob-hole interactions for its cognate effector TM helices, and interacts with TM helices via side-chain to main-chain hydrogen bonds, we conclude that Eag chaperones provide an environment that mimics transmembrane helical packing to stabilize prePAAR effector TMDs in the cytoplasm prior to effector export from the cell.

**Figure 5.**
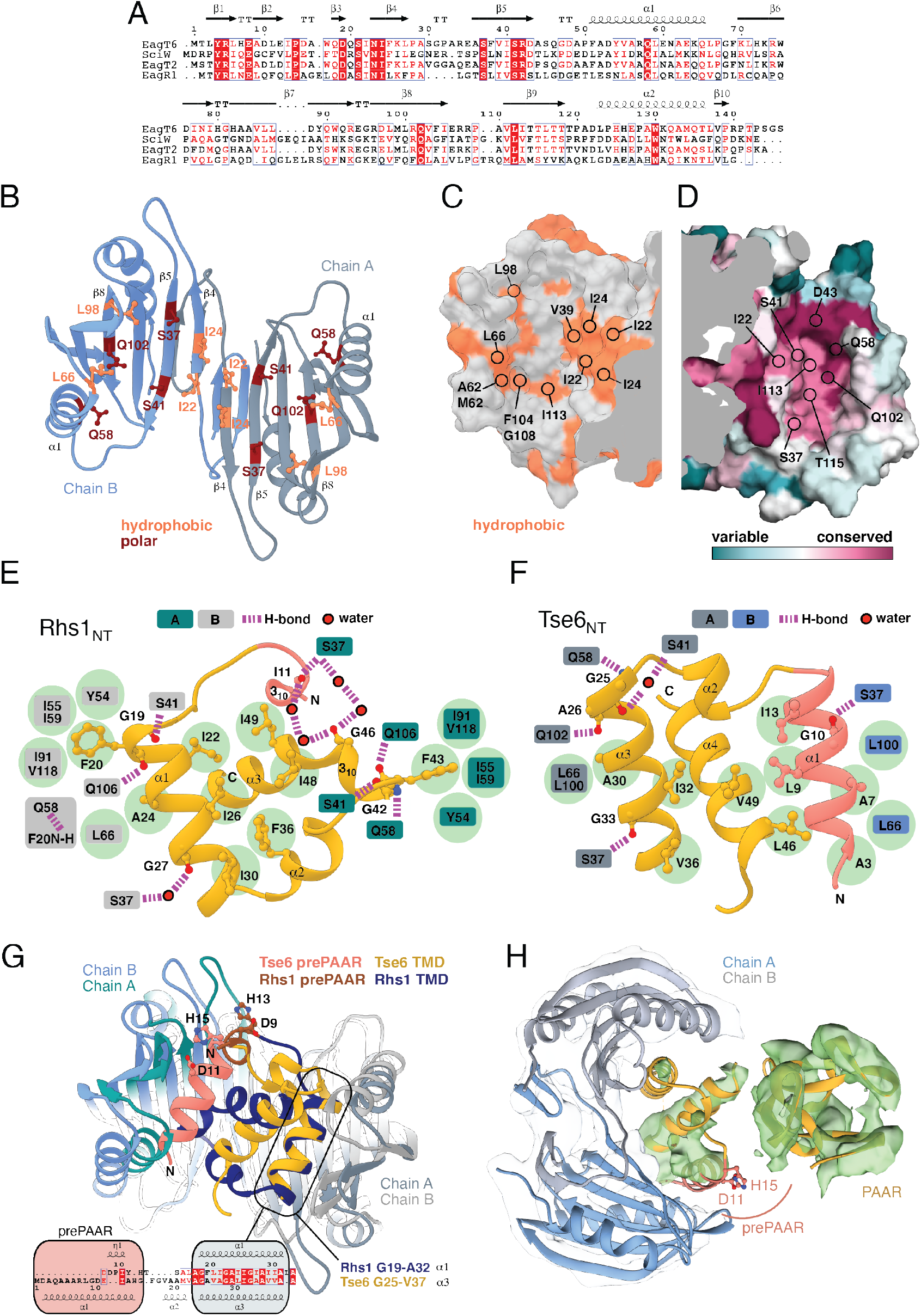
Eag chaperones interact with effector TMDs by mimicking interhelical interactions of alpha helical membrane proteins. A) Alignment of Eag chaperones that interact with class I (SciW, EagR1) or class II (EagT6 and EagT2) prePAAR effectors plotted with secondary structure elements. B) Residues making intimate molecular contacts with their respective TMDs that are conserved among SciW, EagR1, EagT6 and EagT2 are shown. Hydrophobic contacts are colored in light orange and polar contacts in deep red. Residue numbers are based on EagT6. C and D) The conserved hydrophobic molecular surface of the chaperones is shown in light orange (C) and their molecular surface residue conservation is shown as determined by the Consurf server (D)(Ashkenazy et al., 2016). Conserved residues making contacts with the TMDs in both co-crystal structures are shown. E) Molecular contact map of Rhs1_NT_ (residues 1-59) and SciW. prePAAR is shown in pink and the TMD regions in gold. Amino acids making contacts with the conserved residues of the Eag chaperones are shown by side chain/and or by main chain atoms (red for carbonyl and blue for amide). Residues in the Eag chaperone are highlighted by color of chain A or B. Polar (H-bond) contacts are drawn with a purple dashed line and are made with the side chain of the listed Eag residue. Outlined red circles indicate a water molecule. Light green circles indicate van der Waals interactions and hydrophobic interactions. The central group of hydrophobic residues without a listed chaperone residue all pack into the Eag hydrophobic face in Figure 4G (EagT6 I22/24 and V39). F) Molecular contact map of Tse6_NT_ (residues 1-61) and EagT6. Schematic is the same as panel B. Q102 in EagT6 corresponds to Q106 in SciW. G) Structural alignment of SciW-Rhs1_NT_ and EagT6-Tse6_NT_ co-crystal structures using the structurally conserved TM helix as a reference. Eag chain coloring is the same as Figure 4. Rhs1_NT_ is colored in dark blue with a brown prePAAR and Tse6_NT_ in gold with a pink prePAAR. The conserved solvent accessible prePAAR residues D9/11 and H13/15 are shown in ball and stick model. Inset sequence alignment reflects the structurally aligned residues of Rhs1_NT_ (top) and Tse6_NT_ (bottom) as calculated by UCSF Chimera (Pettersen et al., 2004). Secondary structural elements are labeled. H) Docking of the EagT6-TMD crystal structure from Figure 4C into the previously obtained cryo-EM density map of the EagT6-Tse6-EF-Tu-Tsi6-VgrG1a complex (EMD-0135). Cryo-EM density corresponding to EagT6 is depicted in transparent grey and Tse6-TMD and Tse6-PAAR in green; prePAAR residues are shown in pink. Note that Tse6-TMD was docked independent of EagT6 into the Tse6 density. One of three possible orientations for the PAAR domain is shown.

### prePAAR facilitates PAAR domain folding and interaction with the VgrG spike

We next compared the conformation of the bound prePAAR-TMD fragments between our effector-chaperone co-crystal structures. Interestingly, despite the abovementioned similarities between the SciW and EagT6 structures, the conformation of the N-terminal fragment of their bound prePAAR effector differs significantly. In the SciW complex, Rhs1_NT_ adopts an asymmetric binding mode whereby the effector fragment does not make equivalent molecular contacts with both chains of the two-fold symmetrical chaperone dimer (Figures 4A and 5F). The first TM helix (residues 19-33) binds to the hydrophobic cavity of one SciW protomer whereas the remaining hydrophobic region of Rhs1, which consists of two anti-parallel alpha-helices connected by a short 3_10_ helix, occupies the remainder of the binding surface. Phenylalanine residues F20 and F43 likely play an important role in the asymmetric binding of Rhs1 to SciW because their hydrophobic side chains insert into equivalent hydrophobic pockets found in each SciW protomer (Figure 5E). By contrast, Tse6_NT_ exhibits a pseudosymmetric binding mode with EagT6 (Figure 4C and 5F). In this structure, two alpha-helices of Tse6 each occupy equivalent Eag binding pockets and run in the opposite direction to match the antiparallel arrangement of the EagT6 dimer. For example, A7 and A30 of Tse6 interact with equivalent sites in their respective chaperone protomers (Figures 4B and 5E). These two helices, which consist of prePAAR and a TM helix, flank a central TM helix whose C-terminus extends into the solvent, likely indicating the location of the downstream PAAR domain in the full-length effector.

A lack of interpretable electron density prevented modelling of Rhs1’s entire AARxxDxxxH prePAAR motif in our Rhs1_NT_-SciW co-crystal structure. However, the DxxxH portion of this motif is part of a short 3_10_ helix that orients the aspartate and histidine side chains such that they face outward into solvent (Figure 5—figure supplement 1E-G). By contrast, we were able to model the entire prePAAR motif of Tse6_NT_ and in this case, the motif forms an alpha helix that binds the hydrophobic pocket of an EagT6 protomer. In this structure, the two conserved alanine residues of prePAAR make contact with the EagT6 chaperone whereas the arginine, aspartate and histidine residues are solvent exposed (Figure 4C and 5F). Remarkably, despite existing in different secondary structure elements, the D11 and H15 prePAAR residues of Tse6 are located in a similar 3D location as their D9 and H13 counterparts in Rhs1 (Figure 5G). It should be noted that the modelled conformation of Tse6_NT_ appears to be locked into place by crystal packing suggesting that in solution, Tse6’s prePAAR motif may exhibit significant conformational flexibility and can dissociate from EagT6 as is observed for the prePAAR motif of Rhs1 (Figure 5—figure supplement 1H-I). In support of this, we previously showed that addition of detergent disrupts the interaction between EagT6 and Tse6 suggesting that Eag chaperone-effector interactions are labile, likely because chaperone dissociation is required prior to effector delivery into target cells (Quentin et al., 2018). Intriguingly, docking our high resolution EagT6-Tse6_NT_ crystal structure into our previously determined lower resolution Tse6-EagT6-VgrG1 cryo-EM map orients the D11 and H15 prePAAR residues of Tse6 in a position that suggests they interact with its PAAR domain (Figure 5H). In sum, our structural analyses of prePAAR shows that this region is likely dynamic, and its mode of interaction varies for class I and class II prePAAR effectors. However, both Eag chaperones bind the N-terminus of their cognate effector such that the conserved aspartate and histidine residues of prePAAR are positioned to potentially be involved in interactions with the downstream PAAR domain, and thus may play a role in effector-VgrG interactions.

To test if prePAAR influences PAAR function, we next conducted mutagenesis analysis on Tse6 because its PAAR-dependent interaction with its cognate VgrG protein, VgrG1a, can be monitored *in vivo* by western blot. During denaturing electrophoresis, Tse6 appears in two forms: 1) a high-molecular weight species corresponding to Tse6-VgrG1a complex and 2) a low-molecular weight species indicative of free Tse6 (Whitney et al., 2015). Deletion of *vgrG1a* only affects complex formation whereas deletion of the *eagT6* gene results in a substantial reduction in the levels of both species, which provides a means to differentiate residues involved in effector-chaperone versus effector-VgrG interactions (Quentin et al., 2018). Using this readout, we engineered *P. aeruginosa* strains expressing Tse6 D11A and H15A single amino acid substitutions and a D11A/H15A double substitution and examined the consequences of these prePAAR mutations on Tse6 interactions. In support of a role in promoting proper folding of PAAR, Tse6-VgrG1a complex formation was substantially reduced in a strain expressing the Tse6^D11A^ variant and abolished in a strain expressing Tse6^D11A, H15A^ (Figure 6A). We next examined the effect of these mutations on T6SS-dependent delivery of Tse6 into target cells by subjecting these *P. aeruginosa* strains to growth competition assays against Tse6-sensitive recipients. In agreement with our biochemical data, strains expressing Tse6 harboring a D11A mutation exhibited a substantial reduction in co-culture fitness consistent with an inability of these mutant proteins to form a complex with VgrG1a (Fig. 6B).

**Figure 6.**
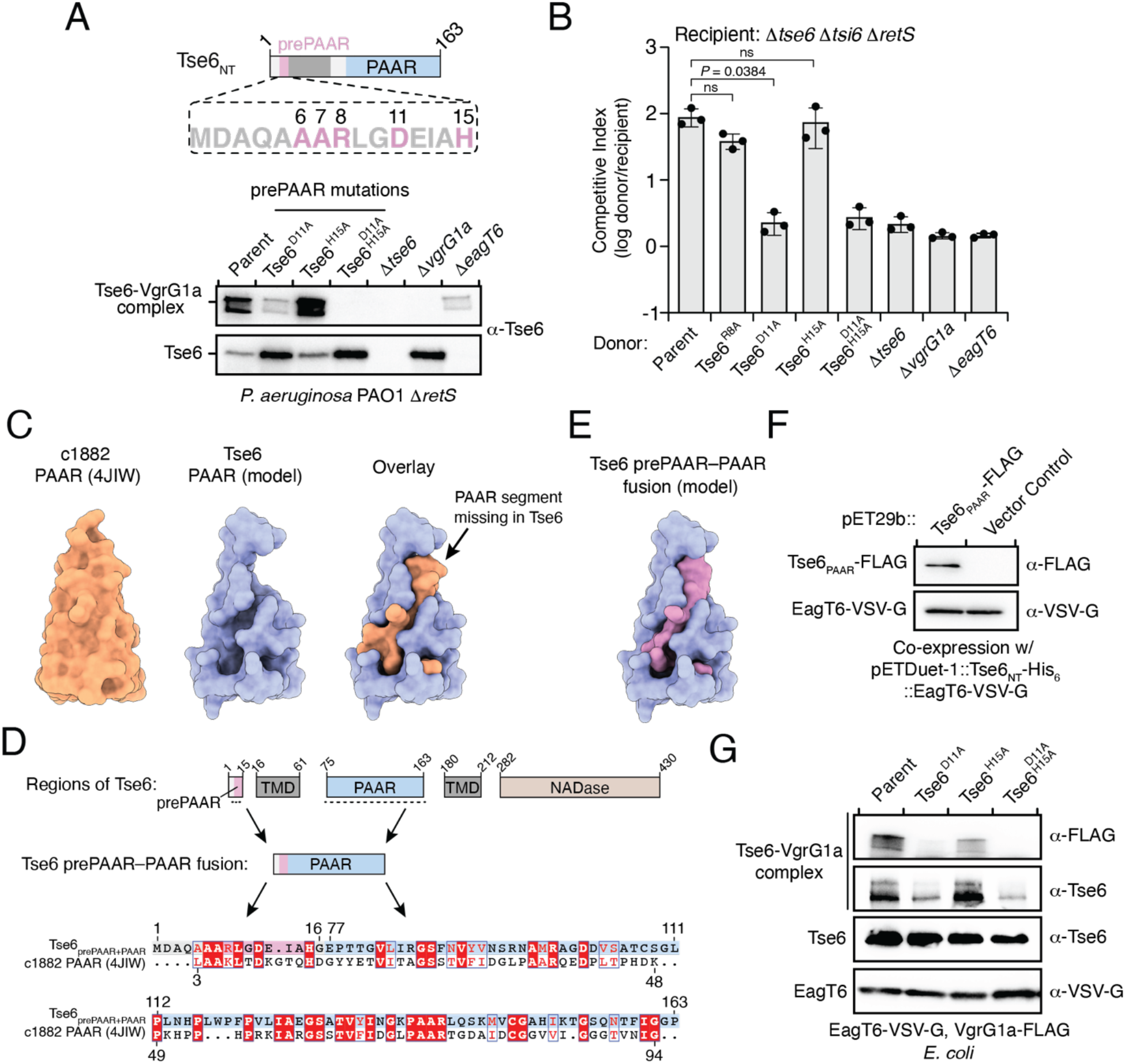
prePAAR is required for PAAR domain interaction with the VgrG spike. **A)** Western blot analysis of Tse6 from cell fractions of the indicated *P. aeruginosa* strains. Low-molecular weight band indicates Tse6 alone whereas high-molecular weight band indicates Tse6-VgrG1a complex. The parental strain contains a Δ*retS* deletion to transcriptionally activate the T6SS (Goodman et al., 2004). B) Outcome of growth competition assay between the indicated *P. aeruginosa* donor and recipient strains. The parent strain is *P. aeruginosa* Δ*retS*. Data are mean ±s.d. for *n* = 3 biological replicates; *P* value shown is from a two-tailed, unpaired *t*-test; ns indicates data that are not significantly different. C) Structural comparison of the c1882 PAAR protein from *E. coli* (PDB: 4JIW) with a model of the PAAR domain of Tse6 generated using Phyre^2^ (Kelley et al., 2015). The overlay shows the additional N-terminal segment present in c1882 that is absent in Tse6. C and D) Schematic showing the residue boundaries of the different regions of Tse6. The prePAAR (pink) and PAAR (blue) sequences were artificially fused to generate Tse6_prePAAR+PAAR_ and used to generate an alignment with c1882 (C) and a structural model (D). Pink space-filling representation indicates the region of the model comprised of prePAAR. F and G) Western blot of elution samples from affinity pull-down of His_6_-tagged Tse6_NT_, containing only prePAAR and the first TMD (residues 1-61), co-expressed in *E. coli* with EagT6 and the PAAR domain of Tse6 with the indicated epitope tags (F) or with the indicated His_6_-tagged Tse6 variants co-purified with VgrG1a-FLAG and EagT6-VSV-G in *E. coli* (G). A, F, G) Data are representative of two independent experiments.

To better understand why Tse6’s PAAR domain requires prePAAR for function, we compared its sequence and predicted structure to the X-ray crystal structure of the ‘orphan’ PAAR domain c1882 from *E. coli*, which does not contain additional components such as TMDs or a toxin domain (Shneider et al., 2013). Interestingly, this analysis suggested that the PAAR domain of Tse6 lacks an N-terminal segment, which, based on the structure of c1882, is likely important for the proper folding of this domain (Figure 6C). We next extended this structural analysis to include all PAAR domains of the prePAAR effectors that we experimentally confirmed bind Eag chaperones. In all cases, the N-terminal segment of each PAAR domain was missing (Figure 6—figure supplement 1A). We also noted that the prePAAR motif possesses significant sequence homology to the N-terminal segment of c1882, suggesting that even though this stretch of amino acids exists on the opposite side of the first TMD region of Tse6, it may comprise the missing segment of Tse6’s PAAR domain (Figure 6D). Lending further support to this hypothesis, when we artificially fused Tse6’s prePAAR motif (residues 1-16) with its PAAR domain (residues 77-163) and generated a structural model. Strikingly, this analysis suggests that the first 16 residues of Tse6 fill the missing structural elements of Tse6’s PAAR domain (Figure 6E). To test if prePAAR interacts with PAAR as this model predicts, we next co-expressed Tse6_NT_-EagT6 complex with the Tse6’s PAAR domain (residues 75-162) and examined the ability of these Tse6 fragments to co-purify with one another. In line with our hypothesis, upon purification of Tse6_NT_-EagT6 complex, Tse6’s PAAR domain was also present (Figure 6F). Taken together with our *in vivo* data, this observation suggests that interaction with prePAAR is critical for the folding and proper function of PAAR. Lending further support to this idea, co-incubation of Tse6, EagT6 and VgrG1a after overexpression in *E. coli* leads to the formation of SDS-resistant Tse6-VgrG1a complexes whereas doing so with a strain expressing Tse6^D11A, H15A^ does not (Figures 6G and Figure 6—figure supplement 1E). Importantly, these mutations do not affect overall levels of Tse6 in cells or affect its ability to bind to EagT6, indicating that these mutations do not have a global destabilizing effect on Tse6 (Figure 6G). Together, these data suggest that the PAAR domains of prePAAR effectors exist as ‘split PAAR’ due to the presence of N-terminal TMDs.

In orphan PAAR proteins, such as c1882, DxxxH motifs are necessary for Zn^2+^-coordination and are therefore necessary for proper folding of this domain (Shneider et al., 2013). In agreement with this precedent, the conserved histidine residue in the DxxxH portion of Tse6’s prePAAR motif is predicted to be in the same 3D position as the first zinc-coordinating histidine residue of c1882 (Figure 6—figure supplement 1B). To extend this comparison further, we conducted *in silico* analyses to examine potential Zn^2+^-binding residues in 564 orphan PAARs and 1,765 prePAAR effectors and found that while orphan PAAR proteins typically contain a total of four histidine and/or cysteine Zn^2+^-coordinating residues, prePAAR effectors only contain three in their PAAR domain with the fourth likely being provided by the prePAAR motif (Figure 6—figure supplement 1C). In support of this prediction, we found that Tse6-VgrG1a complexes formed by the D11A or H15A variants were susceptible to heat treatment under denaturing conditions whereas the wild-type complex remained intact (Figure 6—figure supplement 1D-E). Collectively, our experimental data and informatics analyses indicate that unlike orphan PAAR proteins, which contain all the necessary molecular determinants for proper folding, prePAAR effectors contain inherently unstable PAAR domains that require a prePAAR motif to ensure their proper folding thus enabling their interaction with their cognate VgrG protein (Figure 7).

**Figure 7.**
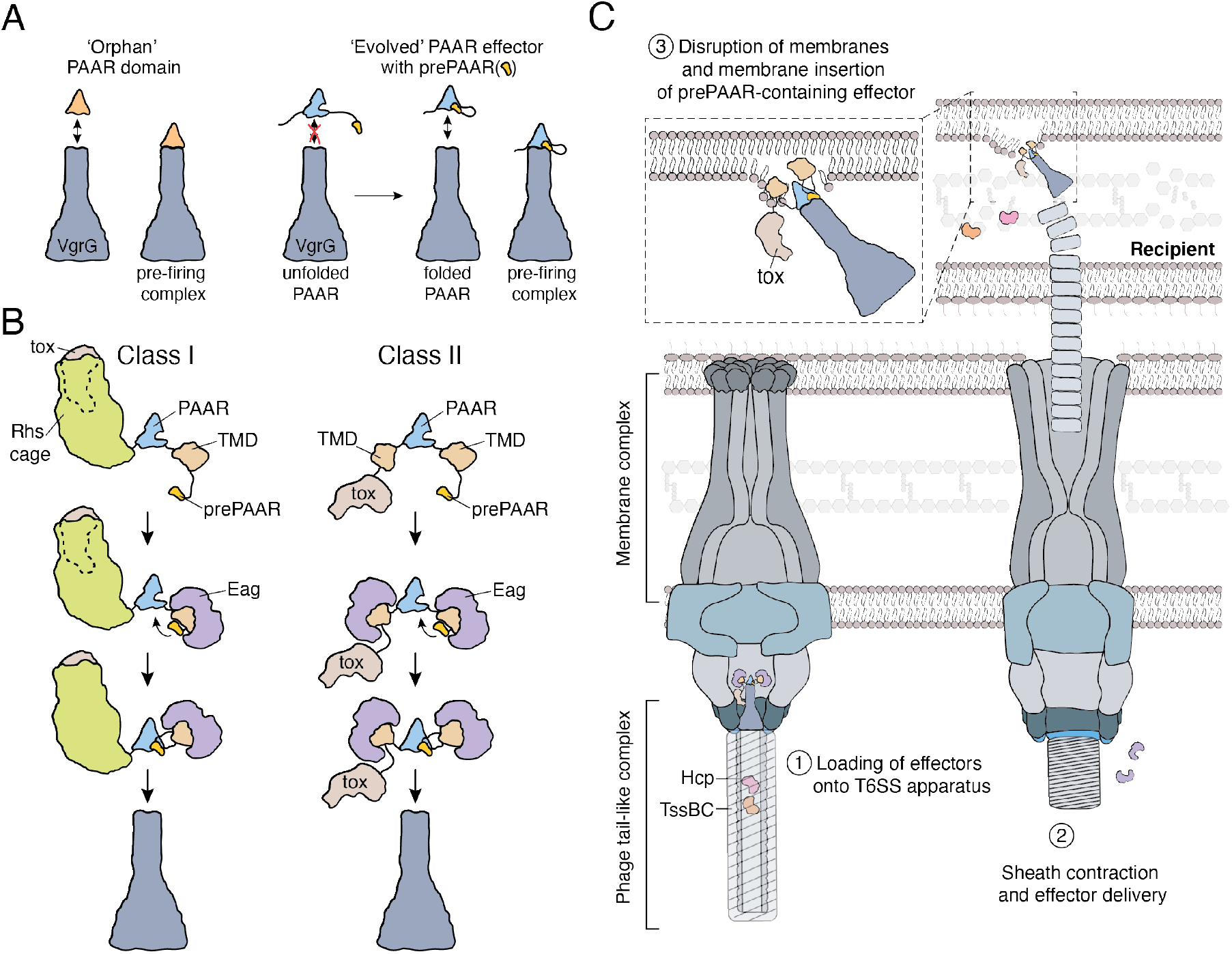
Model depicting the role of Eag chaperones and prePAAR in type VI secretion. A) PAAR proteins exist with or without prePAAR domains. Those that lack prePAAR (orphan), can interact with VgrG and form a functional T6SS spike complex without any additional factors. By contrast, prePAAR-containing effectors contain multiple domains (evolved) and require the prePAAR motif for proper folding of the PAAR domain and thus, loading onto the T6SS apparatus. B) prePAAR-containing effectors can be divided into two classes: class I effectors have a single TMD and contain a C-terminal toxin domain that is likely housed within a Rhs cage whereas class II effectors contain two TMDs and do not possess a Rhs cage. TMD-chaperone and prePAAR-PAAR interactions are required for effector stability and VgrG interaction, respectively, for both classes of prePAAR effectors. C) Depiction of a prePAAR-containing effector being exported by the T6SS into recipient cells. Inset shows the hydrophobic TMDs of a class II prePAAR effector disrupting the inner membrane of the target bacterium to allow entry of the effector toxin domain into the cytoplasm.

## Discussion

Protein secretion systems endow bacteria with a significant fitness advantage in their niche (Galan and Waksman, 2018). The proper functioning of these pathways requires the precise recruitment of effector proteins among hundreds of cytoplasmic proteins. Here, we use a combination of genetic, biochemical and structural approaches to investigate the mechanism of recruitment for a widespread family of membrane protein effectors exported by the T6SS. Our work demonstrates that the N-terminal region of these effectors possesses two structural elements that are critical for their delivery between bacterial cells by the T6SS apparatus. First, this region contains TMDs, which interact with the Eag family of chaperones and are proposed to play a role in effector translocation across the inner membrane of recipient cells (Quentin et al., 2018). Additionally, this region possesses prePAAR, which we show is required for the proper folding of PAAR, thereby facilitating the interaction of this domain with its cognate VgrG protein and enabling effector export by the T6SS.

### The prePAAR motif is present in Eag-binding effectors

prePAAR effectors constitute a new family of T6SS effectors that are defined by the existence of a prePAAR motif, N-terminal TMDs, a PAAR domain and a C-terminal toxin domain. Most notably, we show that this group of effectors co-occurs with Eag chaperones and that chaperone interaction with prePAAR effector TMDs is a conserved property of this protein family. While previous work has relied on genetic context to identify the cognate effector of an Eag chaperone (Whitney et al., 2015, Alcoforado Diniz and Coulthurst, 2015), our use of the prePAAR motif as an effector discovery tool enables the identification of these effectors in any genetic context. Other families of chaperones, such as the DUF4123 or DUF2169 protein families, have also been shown to affect the stability and/or export of their cognate effectors (Burkinshaw et al., 2018, Bondage et al., 2016, Pei et al., 2020). However, little is known about the specificity of these chaperones for their effector targets, which do not contain predicted N-terminal TMDs. DUF4123 chaperones are encoded next to effectors with diverse domain architectures and studies on several members of this family have shown chaperone interactions occur with domains of effectors possessing no apparent shared sequence properties (Liang et al., 2015). A lack of structural information for these and DUF2169 chaperones has hindered an understanding of why certain T6SS effectors require members of these chaperone families for export from the cell.

### Role of Eag chaperones in binding effector TMDs and prePAAR

Our co-crystal structures show that Eag chaperones can exhibit distinct binding modes with the N-termini of their cognate effectors. The class I prePAAR effector Rhs1 interacts with its cognate chaperone SciW in an asymmetric manner whereas the class II effector Tse6 adopts a pseudosymmetric binding mode whereby two separate alpha helices interact with each EagT6 chaperone protomer in a similar location. Our structural analyses suggest that Rhs1 residues F20 and F43 play a critical role in its asymmetric binding mode because the aromatic side chains of these amino acids insert into hydrophobic pockets present in SciW. These favourable chaperone-TMD interactions allow SciW to ‘shield’ the hydrophobic regions of Rhs1’s N-terminus from the aqueous milieu while also positioning its prePAAR motif in such a way that would allow it to interact with PAAR. By contrast, the pseudosymmetric binding mode of Tse6 to EagT6 appears to be much more dynamic as interpretable electron density for bound Tse6 was only observed when the effector fragment was held in place by interactions with an adjacent complex in the crystallographic asymmetric unit. Consequently, we speculate that even though the Tse6’s prePAAR motif appears less accessible than that of Rhs1, it is likely highly dynamic in solution and thus may adopt a markedly different conformation when in complex with PAAR.

Despite containing a primarily beta-sheet secondary structure, Eag chaperones interact with effector TMDs by mimicking the interactions that occur between the helices of alpha-helical membrane proteins, which, to our knowledge, is a unique mechanism for a chaperone-effector interaction. Upon binding their cognate effector, we hypothesize that Eag chaperones not only shield effector TMDs from solvent but also distort their structure to prevent potential hairpin formation and erroneous insertion into the inner membrane of the effector-producing cell. Because Eag-interacting TMDs have likely evolved to insert into bacterial membranes, a mechanism to prevent self-insertion is probably necessary prior to export. Recent work studying the secretion of TMD-containing effectors of the bacterial type III and type IV secretion systems found that shielding TMDs to prevent inner membrane insertion is a critical step for proper targeting to the secretion apparatus (Krampen et al., 2018). However, membrane protein effectors of these secretion systems have evolved to target eukaryotic, not bacterial, membranes and thus may not require stringent control of TMD conformation prior to export. Indeed, unlike the Eag chaperones presented here, a previously studied T3SS chaperone was shown not to distort the conformation of effector TMDs, whose conformation remained similar before and after membrane insertion (Nguyen et al., 2015).

Current evidence also suggests that Eag chaperones are not secreted by the T6SS (Cianfanelli et al., 2016a, Quentin et al., 2018). This leads to two important questions: 1) when do Eag chaperones dissociate from their cognate effector? 2) How do effector TMDs remain stable after their dissociation from the chaperone? Although no definitive answers exist for either of these questions, given that effector-chaperone interactions are maintained after effector-VgrG complex formation, chaperone dissociation presumably occurs immediately before or during a T6SS firing event. One way this could be accomplished is through chaperone interactions with components of the T6SS membrane and/or baseplate subcomplexes, which might induce chaperone-effector dissociation. The lumen of the T6SS apparatus may also serve to mitigate the susceptibility to degradation observed for prePAAR effectors in the absence of Eag chaperones because the inner chamber of the T6SS apparatus may shield effectors from the protein homeostasis machinery of the cell.

### prePAAR-containing proteins contain C-terminal toxin domains that act in the cytoplasm

Studies conducted in several different bacteria suggest that many T6SSs export multiple effectors during a single firing event (Cianfanelli et al., 2016a, Silverman et al., 2013, Hood et al., 2010). The precise subcellular location for effector delivery in recipient cells is not well understood, however, it is noteworthy that many effectors that interact with Hcp or C-terminal extensions of VgrG target periplasmic structures such as peptidoglycan or membranes (Flaugnatti et al., 2016, Silverman et al., 2013, Brooks et al., 2013, LaCourse et al., 2018). In contrast, all characterized prePAAR proteins act on cytoplasmic targets by mechanisms that include the hydrolysis of NAD^+^ and NADP^+^, ADP-ribosylation of FtsZ, pyrophosphorylation of ADP and ATP, and deamination of cytidine bases in double-stranded DNA (Whitney et al., 2015, Ting et al., 2018, Ahmad et al., 2019, Mok et al., 2020). This observation supports the proposal that the TMDs in prePAAR effectors function to promote toxin entry into the cytoplasm of target cells (Quentin et al., 2018). Two possibilities for how this occurs include a discrete toxin translocation event that takes place after the initial delivery of effectors into the target cell periplasm or that effectors are delivered directly into the target cell cytoplasm during a T6SS firing event. The large size of Rhs repeat-containing class I prePAAR effectors favours the latter model because it is unlikely that the 2-3 N-terminal TM helices found in these proteins could form a translocation pore for the C-terminal toxin domain. Instead, we propose that the TMDs of prePAAR effectors acts as molecular grease that coats the tip of the VgrG spike allowing it to effectively penetrate target cell membranes during a T6SS firing event. It should be noted that PAAR effectors with nuclease activity that lack N-terminal TMDs have been identified, suggesting that other cell entry mechanisms likely exist and future work may address whether these proteins have important motifs or domains that permit an alternative translocation mechanism into recipient cells (Pissaridou et al., 2018).

### prePAAR is required for proper PAAR folding and effector export by the T6SS

Crystal structures of single domain PAAR proteins suggest that this domain folds independently and is highly modular (Shneider et al., 2013). Indeed in many instances, PAAR domains appear in isolation (orphan PAAR) and do not require additional binding partners to interact with VgrG (Wood et al., 2019). The initial characterization of PAAR domains established seven groups of PAAR proteins, with the most abundant being orphan PAARs (55% of 1353 PAAR proteins) while the remaining groups represent PAAR proteins with N- and/or C-terminal extensions (Shneider et al., 2013). Our data demonstrate that PAAR domains with N-terminal extensions possess prePAAR, which we show is required for the proper folding of the downstream PAAR domain. Based on our structural modelling and sequence alignments, the ability of prePAAR to assist with PAAR domain folding may in part be due to its participation in coordinating the zinc ion found near the tip of this cone-shaped protein. Our sequence analysis also suggests that while orphan PAARs contain four zinc-coordinating histidine and/or cysteine residues, the PAAR domain of prePAAR effectors contains only three, suggesting that the fourth ligand required for tetrahedrally coordinated Zn^2+^ is provided by prePAAR. In this way, the PAAR domain of prePAAR effectors is split into two components, which come together to form a structure that can interact with VgrG and undergo T6SS-mediated export. One consequence of this ‘split PAAR’ domain arrangement is that the TMDs are tethered to PAAR via their N- and C-terminus, which would restrict the mobility of the TMDs and ensure their positioning on the surface of PAAR. We speculate that the proper arrangement of prePAAR effector TMDs on the surface of PAAR is likely critical for the ability of the T6SS spike complex to effectively penetrate target cell membranes during a T6SS firing event. Future studies focused on capturing high-resolution structural snapshots of assembled prePAAR-TMD-PAAR complexes will be needed to further support this proposed mechanism.

## Conclusions

In summary, our mechanistic dissection of prePAAR effectors and their cognate chaperones has revealed fundamental new insights into bacterial toxin export and membrane protein trafficking. The unique ability of T6SSs to potently target a wide range of bacteria in a contact-dependent manner may permit their use in different biomedical applications, such as the selective depletion of specific bacterial species in complex microbial communities (Ting et al., 2020). An in-depth understanding of the mechanisms that that underlie T6SS effector recruitment and delivery will be of critical importance for such future bioengineering efforts.

## Acknowledgements

The authors would like to thank Jianhua Zhao for electron microscopy expertise, Sarah Trilesky and Matthew Walker for their assistance with cloning and protein purification and Peter Stogios, Seemay Chou, James Holton and Atanas Radkov for crystallography expertise. S.A. and K.K.T. were supported by Ontario Graduate Scholarships and A.G.M. holds a Cisco Research Chair in Bioinformatics. Part of the research described in this paper was performed using beamline 08ID-1 at the Canadian Light Source, a national research facility of the University of Saskatchewan, which is supported by the Canada Foundation for Innovation (CFI), the Natural Sciences and Engineering Research Council (NSERC), the National Research Council (NRC), the Canadian Institutes of Health Research (CIHR), the Government of Saskatchewan, and the University of Saskatchewan. This work was supported by the Max Planck Society (to S.R.) and by grants from CFI (34531 to A.G.M. and 37841 to G.P.), NSERC (RGPIN-2017-05350 to J.C.W. and RGPIN-2018-04968 to G.P.) and CIHR (PJT156129 to J.C.W. and PJT156214 to A.G.M.). Computer resources were supplied by the McMaster Service Lab and Repository computing cluster, funded in part by grants to A.G.M. from CFI and Compute Canada (www.computecanada.ca).

## Author Contributions

Experiments were conceived and designed by S.A., G.P. and J.C.W. All cloning, strain generation, bacterial competition assays and biochemical experiments were conducted by S.A. Bioinformatics analyses were conducted by K.K.T. Protein crystallization, X-ray data collection and analysis was performed by K.S. and G.P. Negative-stain EM experiments were conducted by D.Q. Assistance with cloning and biochemical experiments was provided by T.M.T. Figure design, manuscript writing and editing were done by S.A., G.P., J.C.W. The project was supervised by G.P. and J.C.W. Funding was provided by S.R., A.G.M., G.P. and J.C.W.

## Experimental Procedures

### Bacterial strains and growth conditions

*Pseudomonas* strains used in this study were derived from *P*. *aeruginosa* PAO1 and *P*. *protegens* Pf-5 (Table 5). Both organisms were grown in LB medium (10 g L^−1^ NaCl, 10 g L^−1^ tryptone, and 5 g L^−1^ yeast extract) at 37°C (*P*. *aeruginosa*) or 30°C (*P*. *protegens*). Solid media contained 1.5% or 3% agar. Media were supplemented with gentamicin (30 μg mL^−1^) and IPTG (250 μM) as needed.

**Table 5:**
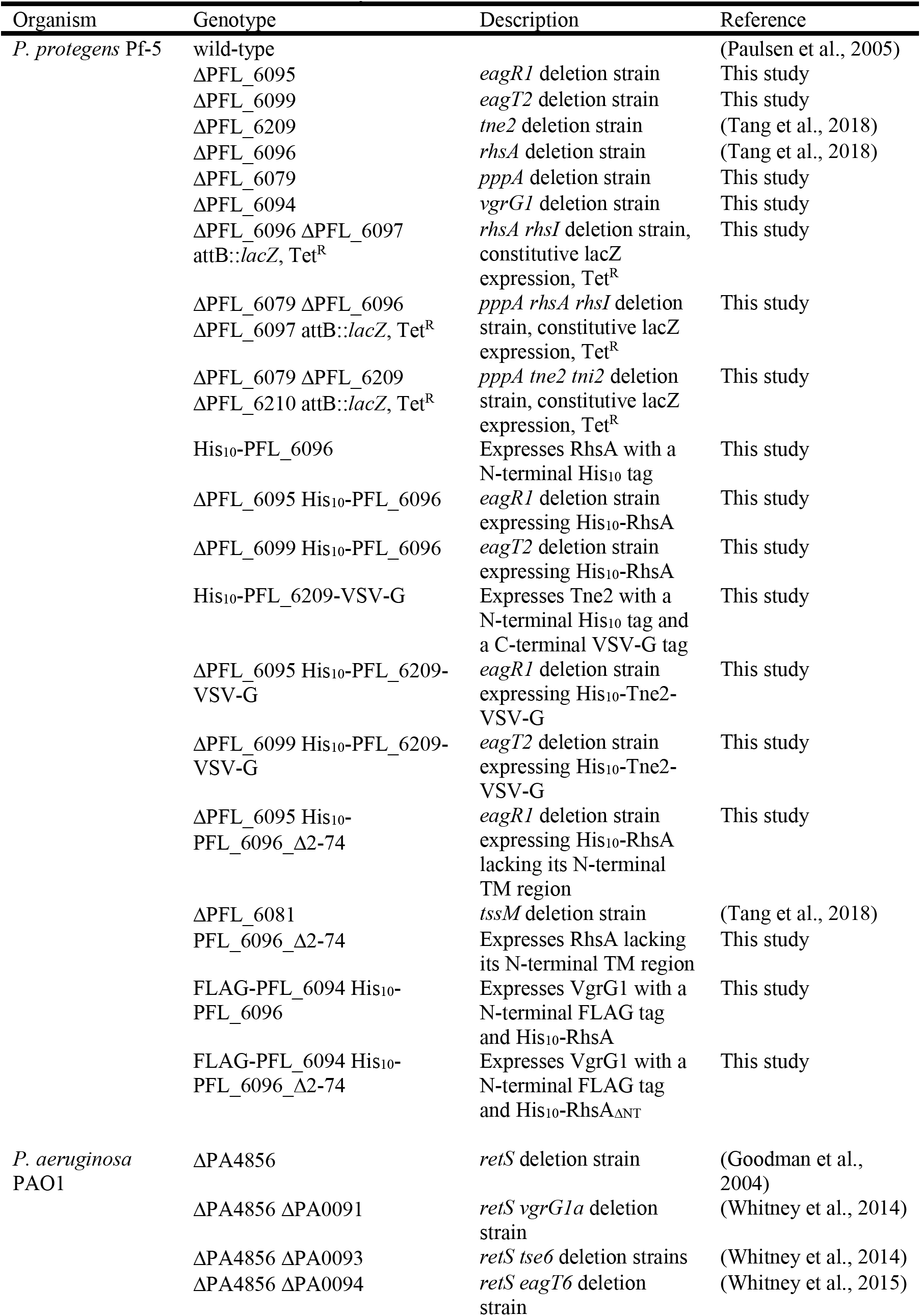
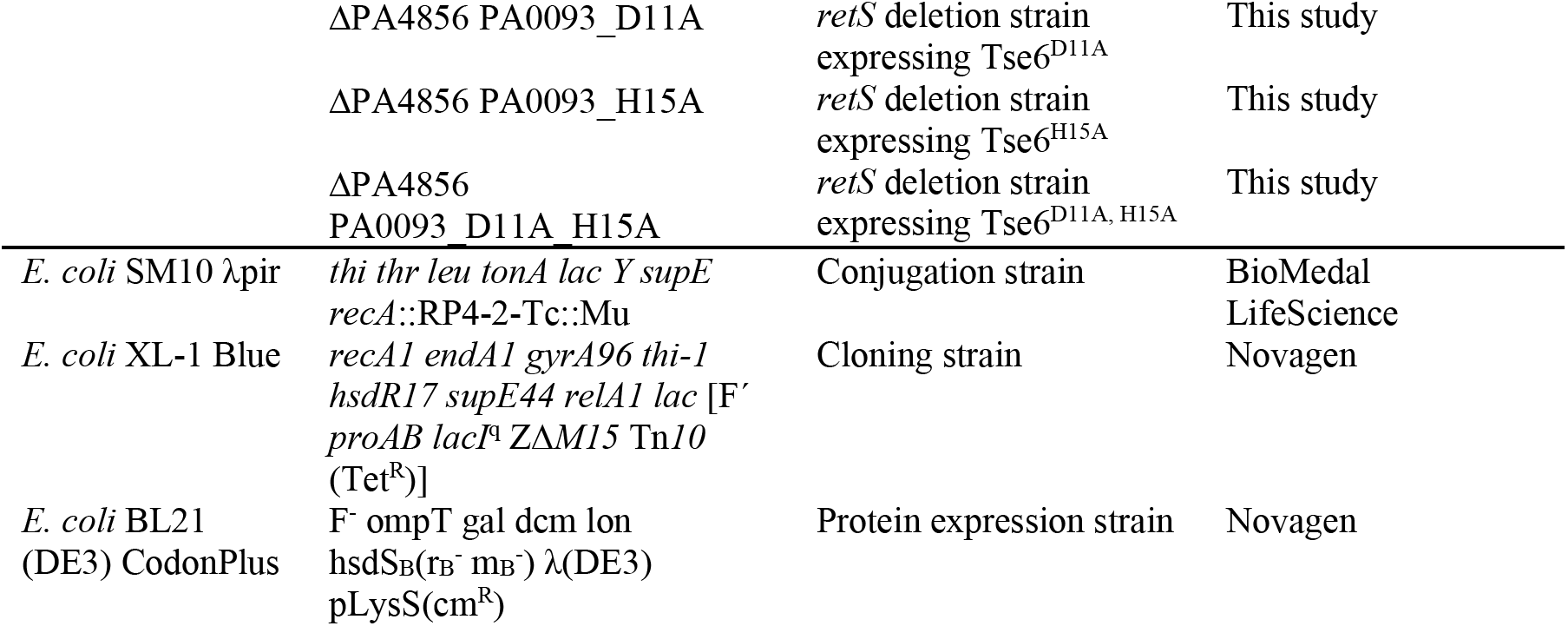
Strains used in this study.

*Escherichia coli* strains XL-1 Blue, SM10 and BL21 (DE3) Gold or CodonPlus were used for plasmid maintenance and toxicity experiments, conjugative transfer and protein overexpression, respectively (Table 5). All *E. coli* strains were grown at 37°C in LB medium. Where appropriate, media were supplemented with 150 μg mL^−1^ carbenicillin, 50 μg mL^−1^ kanamycin, 200 μg mL^−1^ trimethoprim, 15 μg mL^−1^ gentamicin, 0.25-1.0 mM isopropyl β-D-1-thiogalactopyranoside (IPTG), 0.1% (w/v) rhamnose or 40 μg mL^−1^ X-gal.

### Genomic analyses of effector sequences in UniProtKB

For the analysis of all effectors in UniprotKB we used six iterations of *jackhmmer* (HmmerWeb v2.41.1) using the first 60 amino acids of Tse6 (PA0093) protein to obtain 2,378 sequences. We removed any UniProtKB deprecated sequences entries (324/2378, remaining: 2,054) and further filter, cluster, and analyze the remaining 975 effector sequences as stated below (same as analysis in Figure 1E). In our PAAR motif search, using our first to fourth PAAR motif HMMs (see analysis below), we identified 734/975, 200/241, 30/41, and 8/11 sequences to respectively have PAAR motifs. The remaining 3 sequences that did not have PAAR motifs were determined to either directly associated with a PAAR domain downstream. There were 7 sequences that did not have any predicted TM. All scripts and intermediate files can be found in: https://github.com/karatsang/effector_chaperone_T6SS/tree/master/UniProtKB

### Genomic analyses of effector sequences in T6SS-containing genera

The genome assemblies of *Pseudomonas*, *Burkholderia*, *Enterobacter*, *Escherichia*, *Salmonella*, *Serratia*, *Shigella*, *Vibrio* and *Yersinia* were downloaded from NCBI using ncbi-genome-download (https://github.com/kblin/ncbi-genome-download, v0.2.10).

Protein coding genes were predicted using Prodigal (v2.6.3) and the ‘-e 1’ option (Hyatt et al., 2010). We developed a Hidden Markov Model (HMM) for detecting effectors by using the first 61 amino acids of Tse6 (PA0093) protein and six iterations of *jackhmmer* (HmmerWeb v2.41.1). *hmmsearch* (v3.1b2) and the effector HMM were used to identify the effectors in all genome assemblies using the ‘ -Z 45638612 -E 1000’ options and we further filtered for a bitscore greater than 40. We further filtered to include effectors that included the prePAAR (AARxxDxxxH) motif, which we searched for using regular expressions, identifying 6,129 prePAAR-containing sequences across 5,584 genomes. To be included in the analysis of Figure 1D, each genome with at least one effector had to also encode for an Eag chaperone which we searched for using Pfam’s established DcrB HMM (http://pfam.xfam.org/family/PF08786#tabview=tab6) and hmmsearch with the same parameter and bitscore cutoff as the effector search. For Figure 1E, to reduce spurious effector predictions, we removed sequences with less than 100 amino acids. To reduce redundancy, we removed any sequences that were 100% identical and clustered sequences with 95% sequence similarity that were less than 50 amino acids different in length using CD-HIT (v4.8.1 with ‘ -c 0.95 -n 5 -S 50’), leaving 1,166 sequences for further analysis (Li and Godzik, 2006). The numbers of sequences before and after filtering for the UniprotKB and sequences isolated from the 8 genera listed above are indicated in Table 3. We identified the presence of a PAAR domain through a repetitive process of generating a PAAR motif HMMs and using *hmmsearch* (as described above) to capture the known diversity of the PAAR motif. We started broadly by using Pfam’s PAAR motif HMM (http://pfam.xfam.org/family/PF05488#tabview=tab4) to identify 895/1166 PAAR motif containing sequences. For the 271 sequences that were predicted to not have a PAAR motif, we then generated an HMM using three iterations *jackhmmer* and the PAAR motif of the Tse6 (PA0093) protein (L75 to G162) to identify 219/271 PAAR motifs. We generated a third PAAR motif HMM using 60-160 amino acids of a randomly selected sequence (GCF_001214785.1 in contig NZ_CTBP01000066.1) and two iterations of *jackhmmer* that was not identified to have a PAAR motif in the previous search but was identified to have a PAAR domain using phmmer (HmmerWeb version 2.41.1). We identified 42/52 sequences had a PAAR domain using the third PAAR motif. For the fourth PAAR domain HMM, we used the 60-160 amino acid sequence of GCF_005396085.1 in the NZ_BGGV01000116 contig and three iterations of *jackhammer* to identify 8/10 sequences that had a PAAR motif. The remaining two sequences with no PAAR domain were manually analyzed and were determined to either be directly associated with a PAAR domain downstream (GCF_001425105.1) or directly beside T6SS machinery gene (GCF_001034685.1). We predicted the transmembrane (TM) helices in proteins first using TMHMM (v2.0), Phobius web server, and TMbase (https://embnet.vital-it.ch/software/) (Krogh et al., 2001, Kall et al., 2007). Using TMHMM, we defined a TM region to include TM helices that were less than or equal to 25 amino acids apart. Therefore, any TM helix that was greater than 25 amino acids apart from another TM helix would be considered part of a new TM region. Any effector considered to have no TM or three TM regions were analyzed with Phobius with the same criteria as with TMHMM. Any effector considered to have no TM or three TM regions using Phobius, were analyzed with TMbase where we used the strongly preferred model and only interpreted TM helices with a score greater than 1450. In this model, any TM helices within the first 120 amino acids is one TM region and any number of TM helices between 200 and 300 amino acids were another region. MAFFT (v7.455) was used to align the sequences using the ‘--auto’ option and the alignment was then trimmed to remove gaps using trimAl (v1.4) and the ‘-gt 0.8 -cons 80’ options (Katoh and Standley, 2013, Capella-Gutierrez et al., 2009). We constructed the maximum-likelihood phylogenetic tree using FastTree (v2.1.10) and the ‘-gamma’ option(Price et al., 2010). The phylogenetic tree was visualized using ggtree (Yu, 2020). For Figure 1—figure supplement 1B, we identified neighbouring (within 300 base pairs) chaperone sequences for the effectors in Figure 1E. We removed any effectors that did not have a chaperone and we categorized the chaperones with its corresponding effectors TM prediction. Sequence logos in Figure 1C and 1F were created by using logo maker (v0.8) (Tareen and Kinney, 2020). All scripts and intermediate files can be found in: https://github.com/karatsang/effector_chaperone_T6SS/tree/master/NCBI_8_Genera.

### Screening for potential Zn^2+^-binding residues

To collect orphan PAAR sequences, we used the Pfam database’s information on the PAAR motif (PF05488, http://pfam.xfam.org/family/PF05488#tabview=tab1) and only obtained the 1,923 sequences with one PAAR motif architecture. We then aligned and trimmed the alignment of the 1,923 orphan PAAR sequences. We then used the previously mentioned 2,054 effector sequences from UniProtKB and filtered to only use 1765 sequences with an AARxxDxxxH motif. To identify Zn^2+^-binding residues in orphan and prePAAR effector sequence logos, we used logo maker (v0.8) to create sequence logos for the first 200 amino acids (Tareen and Kinney, 2020). All scripts and intermediate files can be found in: https://github.com/karatsang/effector_chaperone_T6SS/tree/master/ZnBindingResidues

### DNA manipulation and plasmid construction

Primers were synthesized and purified by Integrated DNA Technologies (IDT). Phusion polymerase, restriction enzymes and T4 DNA ligase were obtained from New England Biolabs (NEB). Sanger sequencing was performed by Genewiz Incorporated.

Plasmids used for heterologous expression were pETDuet-1, pET29b and pSCrhaB2-CV. Mutant constructs were made using splicing by overlap-extension PCR and standard restriction enzyme-based cloning procedures were subsequently used to ligate PCR products into the plasmid of interest.

In-frame chromosomal deletion mutants in *P. aeruginosa* and *P. protegens* were made using the pEXG2 plasmid as described previously (Hmelo et al., 2015). Briefly, 500-600 bp upstream and downstream of target gene were amplified by standard PCR and spliced together by overlap-extension PCR. The resulting DNA fragment was ligated into the pEXG2 allelic exchange vector using standard cloning procedures (Table 6). Deletion constructs were transformed into *E. coli* SM10 and subsequently introduced into *P. aeruginosa* or *P. protegens* via conjugal transfer. Merodiploids were directly plated on LB (lacking NaCl) containing 5% (w/v) sucrose for *sacB*-based counter-selection. Deletions were confirmed by colony PCR in strains that were resistant to sucrose, but sensitive to gentamicin. Chromosomal point mutations or tags were constructed similarly with the constructs harboring the mutation or tag cloned into pEXG2. Sucrose-resistant and gentamicin-sensitive colonies were confirmed to have the mutations of interest by Sanger sequencing of appropriate PCR amplicons.

**Table 6:**
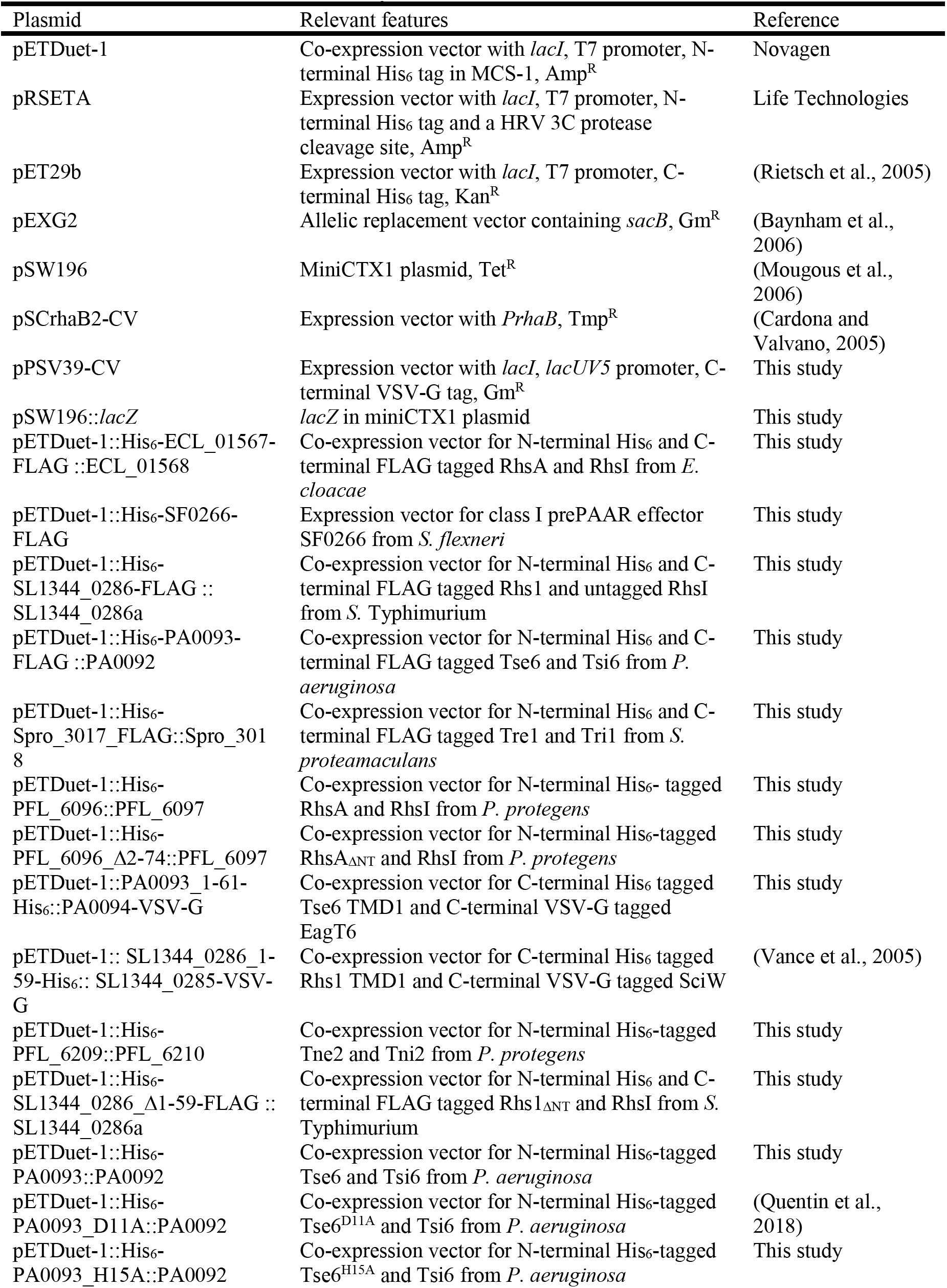
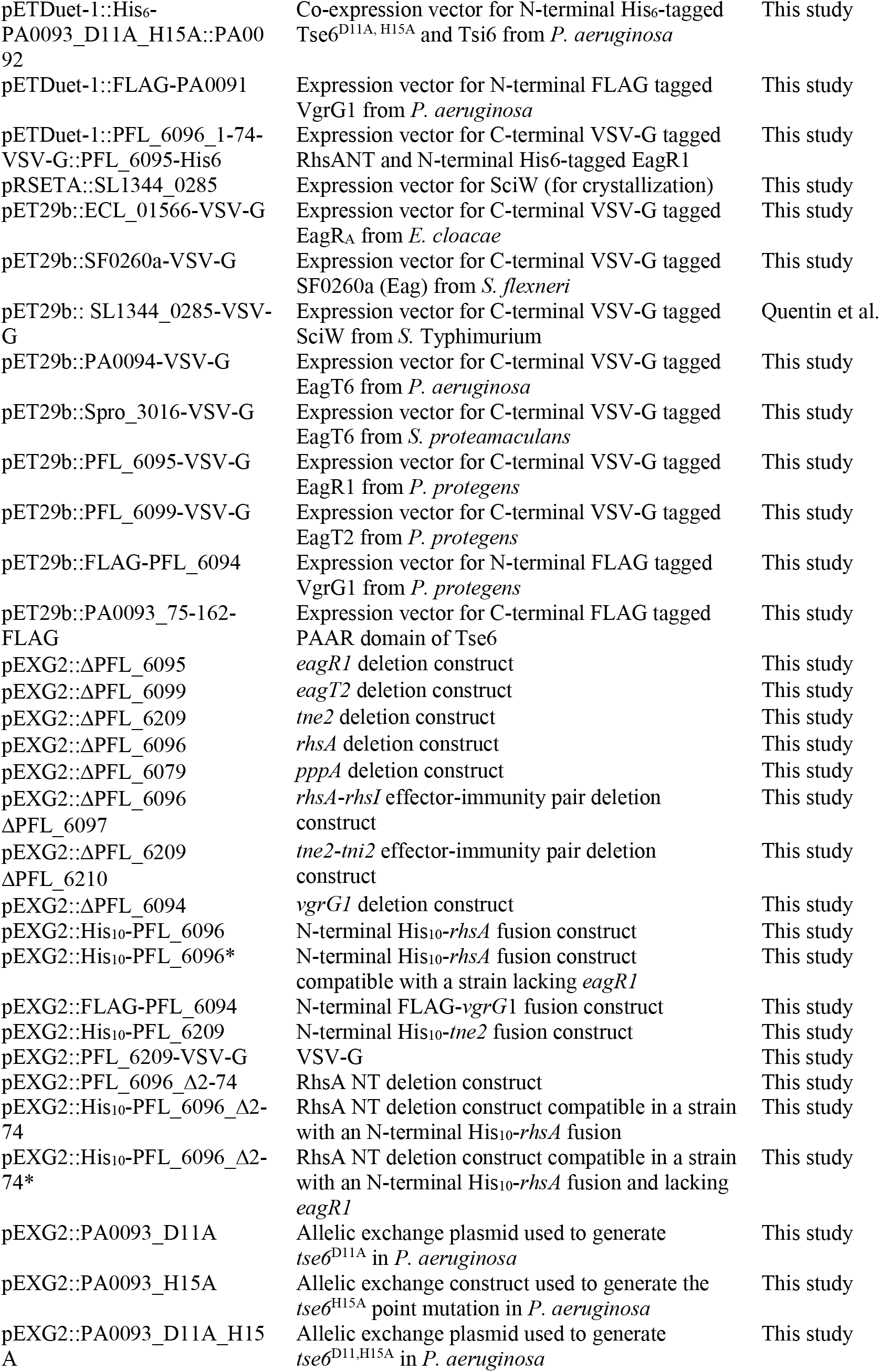
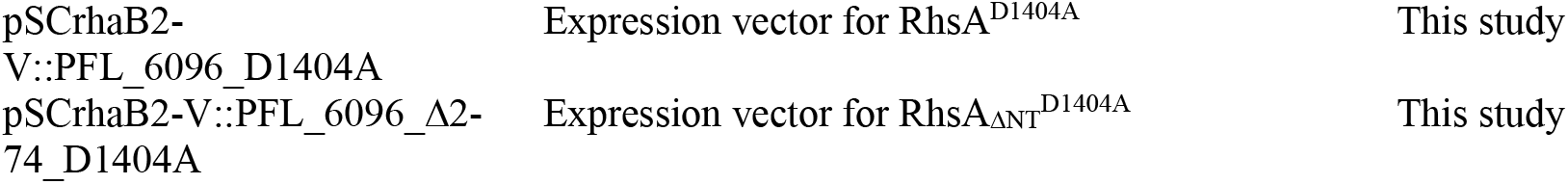
Plasmids used in this study.

### Bacterial toxicity experiments

We previously showed that a D1404A mutation was sufficient to attenuate, but not abolish, the toxicity of RhsA and allows for the cloning of this toxin in the absence of its immunity gene (Tang et al., 2018). Therefore, to assess the toxicity of the full-length effector and a truncated variant, we cloned RhsA^D1404A^ or RhsA_ΔNT_D1404A into the rhamnose-inducible pSCrhaB2-CV vector. These plasmids were co-transformed with an IPTG-inducible pPSV39 vector containing or lacking EagR1, respectively (see Table 6). Stationary-phase overnight cultures containing these plasmids were serially diluted 10^−6^ in 10-fold increments and each dilution was spotted onto LB agar plates containing 0.1% (w/v) L-rhamnose, 250 μM IPTG, trimethoprim 250 μg mL^−1^ and 15 μg mL^−1^ gentamicin. Photographs were taken after overnight growth at 37°C.

### Cell fraction preparation and secretion assays

Stationary-phase overnight cultures of *E. coli* (DE3) BL21 CodonPlus, *P. aeruginosa* Δ*retS* or *P. protegens* were inoculated into 2 mL or 50 mL LB at a ratio of 1:500, respectively. Cultures were grown at 37 °C (*E. coli* and *P. aerugionsa*) or 30 °C (*P. protegens*) to OD 0.6-0.8. Upon reaching the desired OD, all samples were centrifuged at 7, 600 x *g* for 3 min. The secreted fraction in *P. aeruginosa* or *P. protegens* samples was prepared by isolating the supernatant and treating it with TCA (final conc: 10% (v/v)) as described previously (Quentin et al., 2018). The cell pellet was resuspended in 60 μL PBS, treated with 4X laemmli SDS loading dye and subjected to boiling to denature and lyse cells. For experiments examining the stability of Tse6-VgrG1a complexes, *P. aeruginosa* cells were resuspended in 60 μL PBS and subjected to six freeze-thaw cycles prior to mixing with 2X laemmli SDS loading dye. For preparation of *P. protegens* and *E. coli* cell fractions containing his-tagged complexes, cells were resuspended in lysis buffer containing 50 mM Tris-HCl (pH 8.0), 250 mM NaCl, 10 mM imidazole and purified according to the protocol described below (see *Protein expression and purification*).

### Competition assays

A tetracycline-resistant, *lacZ*-expression cassette was inserted into a neutral phage attachment site (*attB*) of recipient *P. aeruginosa* and *P. protegens* strains to differentiate these strains from unlabeled donors. *P. protegens* recipient strains also contain a Δ*pppA* mutation to stimulate T6SS effector secretion to induce a T6SS ‘counterattack’ from *P. protegens* donor strains (Basler et al., 2013).

For intraspecific competitions between *P. aeruginosa* or *P. protegens* donors against isogenic recipient that lack the indicated effector-immunity pairs, stationary-phase overnight cultures were mixed in a 1:1 (v/v) ratio.

Initial ratios of donors:recipients were counted by plating part of the competition mixtures on LB agar containing 40 μg mL^−1^ X-gal. The remainder of each competition mixture was spotted (10 μL per spot) in triplicate on a 0.45 μm nitrocellulose membrane overlaid on a 3% LB agar plate and incubated face up at 37 °C for 20-24 h. Competitions were then harvested by resuspending cells in LB and counting colony forming units by plating on LB agar containing 40 μg mL^−1^ X-gal. The final ratio of donor:recipient colony forming units were normalized to the initial ratios of donor and recipient strains.

### Protein expression and purification

All plasmids used for co-purification experiments (chaperone-effector pairs, tagged variants of *P. protegens* proteins and Tse6 prePAAR mutants), RhsA-RhsI-EagR1-VgrG complex for negative-stain EM, Hcp (PFL_6089) and RhsA_ΔNT_ used for antibody development or the SciW, EagT6-Tse6_NT_ complex and the SciW-Rhs1_NT_ complex used for crystallization were expressed in *E. coli* BL21 (DE3) Gold or CodonPlus cells. Important differences in expression strategy used are indicated below.

#### Co-purification experiments, preparation of negative stain EM samples, and preparation of samples for antibody development

Chaperone-effector pairs (e, effector; c, chaperone) from: *Pseudomonas aeruginosa* (e: PA0093, c: PA0094), *Salmonella* Typhimurium (e: SL1344_0286, c: SL1344_0285), *Shigella flexneri* (e: SF0266, c: SF3490), *Enterobacter cloacae* (e: ECL_01567, c: ECL_01566) and *Serratia proteamaculans* (e: Spro_3017, c: Spro_3016) were co-expressed using pET29b containing the predicted chaperone and pETDuet-1 harboring the full-length effector and its predicted immunity determinant. A similar co-expression strategy was employed for the RhsA_ΔNT_-RhsI complex, RhsA-RhsI-EagR1-VgrG1 complex, Tse6 and the Tse6 prePAAR variants, Tsi6 and EagT6 (see Table 6 for details). VgrG1a was expressed in isolation in pETDuet-1 and Hcp (PFL_6089) was expressed in pET29b. For *P. protegens*, all purified proteins were expressed from their native locus.

For the expression of chaperone-effector pairs and the Tse6 prePAAR mutants, a 1 mL overnight culture of expression strains was diluted in 50 mL of LB broth and grown at 37°C (*E. coli*) until OD 0.6-0.8. 40 mL overnight cultures were grown for all other of expression strains and were diluted into 2 L of LB broth and grown to OD_600_ 0.6-0.8 in a shaking incubator at 37°C. For most samples, protein expression was induced by the addition of 1 mM IPTG and cells were further incubated for 4.5 h at 37°C. Expression of large protein complexes (>150 kDa) in *E. coli*, such as the chaperone-effector pairs from *Salmonella* and *Enterobacter,* RhsA_ΔNT_-RhsI and RhsA-RhsI-EagR1-VgrG1 complexes were induced at 18 °C and incubated at this temperature overnight. One millilitre overnight cultures of *P. protegens* strains expressing the desired tagged protein was diluted in 50 mL of LB broth and grown at 30°C (*P. protegens*) unitl OD 0.8. Cells were harvested by centrifugation at 9,800 *g* for 10 min following incubation. For the RhsA-EagR1-VgrG1 complex and the experiments containing Tse6 prePAAR mutants, the pellets for cells expressing the cognate VgrG were combined with the pellets containing effectors, as described above. Pellets from 50 mL culture were resuspended in 3.5 mL lysis buffer (50 mM Tris-HCl pH 8.0, 300 mM NaCl, 10 mM imidazole), whereas those from 2 L of culture were resuspended in 25 mL of lysis buffer prior to rupture by sonication (6 x 30 second pulses, amplitude 30%). Cell lysates were cleared by centrifugation at 39,000 *g* for 60 min and the soluble fraction was loaded onto a gravity flow Ni-NTA column that had been equilibrated in lysis buffer. Ni-NTA-bound complexes were washed twice with 25 mL of lysis buffer followed by elution in 10 mL of lysis buffer containing 400 mM imidazole. The Ni-NTA purified complex was further purified by gel filtration using a HiLoad 16/600 Superdex 200 column equilibrated in 20 mM Tris-HCl pH 8.0 150 mM NaCl or phosphate buffered saline (for samples used for antibody development only).

#### Preparation of samples for crystallization

*sciW* (SL1344_0285) was synthesized with codon optimization for *E. coli* and cloned into the vector pRSETA with the restriction sites NdeI/HindIII (Life Technologies). This construct includes an N-terminal 6-his tag and an HRV 3C protease cleavage site (MGSSHHHHHHSSDLEVLFQGPLS). SciW-Rhs1_NT_ and EagT6-Tse6_NT_ complexes were co-expressed using pETDUET-1. Note that the EagT6 construct has a C-terminal VSV-G tag (see Table 6). Cells were grown in LB broth to OD_600_ 0.6 at 37°C at which point protein expression was induced by the addition of 1mM IPTG. The temperature was reduced to 20°C and cultures were allowed to grow overnight. Cells were harvested by centrifugation and resuspended in lysis buffer followed by lysis with an Emulsiflex-C3 (Avestin). The lysate was cleared by centrifugation at 16,000 rpm for 30 minutes and the supernatant passed over a nickel NTA gravity column (Goldbio) followed by washing with 50 column volumes of chilled lysis including PMSF, DNase I, and MgCl_2_. Proteins were eluted with 5 column volumes elution buffer then purified by gel-filtration using an SD75 16/60 Superdex gel filtration column equilibrated in gel-filtration buffer (GF) with an AKTA pure (GE Healthcare). For SciW, after affinity purification the protein was dialyzed in GF buffer O/N at 4°C and the His-tag removed during dialysis using HRV 3C protease. The digested SciW was passed over a nickel NTA gravity column and the flow through was collected. SciW was further purified using an SD75 16/60 Superdex gel filtration column equilibrated in GF buffer

The buffers used were as follows: SciW lysis buffer (20mM TRIS pH 7.5, 500mM NaCl, 20mM imidazole); SciW elution buffer (20mM TRIS pH 7.5, 500mM NaCl, 500mM imidazole); SciW GF buffer (20 mM TRIS pH 7.5, 250mM NaCl, 1mM 2-Mercaptoethanol); SciW-Rhs1_NT_ and EagT6-Tse6_NT_ complexes lysis buffer (20 mM TRIS pH 8.0, 150 mM, 25 mM imidazole); elution buffer (20 mM TRIS pH 8.0, 150 mM, 500 mM imidazole); and GF buffer (20 mM TRIS pH 8.0, 150 mM NaCl, 1mM 2-Mercaptoethanol).

### Crystallization and structure determination

SciW was concentrated to 7, 14 and 22 mg/mL for initial screening using commercially available screens (Qiagen) by sitting-drop vapor diffusion using a Crystal Gryphon robot (Art Robbins Instruments). The crystallization conditions for SciW were 22 mg/mL with a 1:1 mixture of 0.1 M Tris HCL pH 8.5, 25% (v/v) PEG 550 MME at 4°C. EagT6-Tse6_NT_ complex was concentrated to 5, 10 and 20 mg/mL and screened for crystallization conditions as per SciW. The final crystallization conditions were 20 mg/mL with a 1:1 mixture of 0.2M Magnesium chloride, 0.1M Bis-TRIS pH 5.5, and 25% (w/v) PEG 3350 at 4°C. SciW-Rhs1_NT_ complex was concentrated to 15, 20 and 25mg/mL and screened for crystallization as per SciW. The crystallization conditions were 25 mg/mL protein with a 1:1 mixture of 0.2M Ammonium sulfate, 0.1M Bis-TRIS pH 5.5, and 25% (w/v) PEG 3350 at 4°C.

Diffraction data from crystals of SciW and EagT6-Tse6_NT_ complex were collected in-house at 93K using a MicroMax-007 HF X-ray source and R-axis 4++ detector (Rigaku). Diffraction data from SciW-Rhs1_NT_ crystals were collected at the Canadian Light Source at the Canadian Macromolecular Crystallography Facility Beam line CMCF-ID (08ID-1). SciW crystals were prepared by cryo-protection in mother liquor plus 38% PEG 550 MME and flash freezing in liquid nitrogen. Crystals of EagT6-Tse6_NT_ and SciW-Rhs1_NT_ complexes were prepared in the same manner with increasing the concentration of PEG3350 to 35-38%. All diffraction data were processed using XDS (Kabsch, 2010). Phases for SciW were determined by the molecular replacement-single anomalous diffraction (MR-SAD) technique. A home-source data set was collected from SciW crystals soaked in cryo-protectant containing 350 mM NaI for one-minute before flash freezing. EagT6 (PDB: 1TU1) was used as a search model and phases were improved by SAD using the Phenix package (Adams et al., 2010). Phases for both the EagT6-Tse6_NT_ and SciW-Rhs1_NT_ complexes were obtained by molecular replacement using EagT6 (PDB: 1TU1) and SciW as search models, respectively, with the Phenix package. Initial models were built and refined using Coot, Refmac and the CCP4 suite of programs, Phenix, and TLS refinement (Emsley et al., 2010, Murshudov et al., 1997, Winn et al., 2011, Winn et al., 2001). Data statistics and PDB codes are listed in Table 4. The coordinates and structure factors have been deposited in the Protein data Bank, Research Collaboratory for Structural Bioinformatics, Rutgers University, New Brunswick, NY (www.rcsb.org). Molecular graphics and analysis were performed using Pymol (Schrödinger, LLC) and UCSF Chimera (Pettersen et al., 2004).

### Electron microscopy and image analysis

#### Negative stain sample preparation

Four microlitres of each protein sample at a concentration of approx. 0.01 mg/mL was applied onto glow-discharged carbon-coated copper grids. After 45 s of incubation at room temperature, excess liquid was blotted away using Whatman No. 4 filter paper, followed by two washing steps with GF buffer. Samples were then stained with 1 % (w/v) uranyl formate solution and grids stored at RT until usage.

#### Data collection and image analysis

Images were recorded manually with a JEOL JEM-1400 microscope, equipped with a LaB6 cathode and 4k x 4k CMOS detector F416 (TVIPS), operating at 120 kV. For VgrG1, RhsA_ΔNT_, the EagR1-RhsA complex and EagR1-RhsA-VgrG1 complex, a total of 99, 140, 100 and 120 micrographs, respectively, were collected with a pixel size of 2.26 Å. Particles for the VgrG1, RhsA_ΔNT_, EagR1-RhsA complex and EagR1-RhsA-VgrG1 complex datasets were selected automatically with crYOLO using individually pre-trained models, resulting in 18676, 23907, 32078 and 31409 particles, respectively (Wagner et al., 2019). Subsequent image processing was performed with the SPHIRE software package (Moriya et al., 2017). Particles were then windowed to a final box size of 240 x 240 pixel. Reference-free 2-D classification was calculated using the iterative stable alignment and clustering algorithm (ISAC) implemented in SPHIRE, resulting in 2-D class averages of the respective datasets (Yang et al., 2012). Distance measurement were performed with the e2display functionality in EMAN2 (Tang et al., 2007). The placement of the crystal structure into the electron density map (EMD-0135) was done using rigid-body fitting in Chimera (Pettersen et al., 2004). Here, Tse6-TMD and EagT6 of the EagT6-TMD crystal structure were fitted independently as rigid bodies to better describe the density. Due to the distinct shape of the PAAR domain, three different orientations were possible in the docking step, each rotated by 120°. Docking of Tse6-TMD into the density embraced by the second EagT6 described this density less well.

### Western blot analyses

Western blot analyses of protein samples were performed as described previously for rabbit anti-Tse1 (diluted 1:5,000; Genscript), rabbit anti-FLAG (diluted 1:5,000; Sigma), rabbit anti-VSV-G (diluted 1:5,000; Sigma), rabbit anti-Hcp1 (*P. aeruginosa*) (diluted 1:5,000, Genscript) and detected with anti-rabbit horseradish peroxidase-conjugated secondary antibodies (diluted 1:5,000; Sigma) (Ahmad et al., 2019). Rabbit anti-Hcp (*P. protegens*) was used at a 1:5000 dilution. Western blots were developed using chemiluminescent substrate (Clarity Max, Bio-Rad) and imaged with the ChemiDoc Imaging System (Bio-Rad).

## Data Availability

All data supporting the findings of this study are available within the manuscript and its associated supplementary information. X-ray crystallographic coordinates and structure factor files are available from the PDB: SciW (PDB 6XRB), SciW-Rhs1_NT_ (PDB 6XRR), EagT6-Tse6_NT_ (PDB 6XRF). Tables containing all prePAAR effector sequences can be found in Tables 1 and 2.

---

**Table 1. List of prePAAR motif-containing proteins identified in the UniProtKB Database (provided as Table_S1_UniprotKB_prePAAR_D01.xlsx file)**. The document contains two separate sheets. Dataset A corresponds to 2,054 prePAAR-containing sequences that were identified through an iterative search of the UniprotKB using Tse6_NT_. Dataset B corresponds to 975 sequences collected following filtering of dataset A (see methods for details).

**Table 2. List of prePAAR motif-containing proteins from assembled genomes of all species belonging to the genera *Burkholderia*, *Escherichia*, *Enterobacter*, *Pseudomonas, Salmonella*, *Serratia, Shigella* and *Yersinia* (provided as Table_S2_8_genera_prePAAR_D01.xlsx file)**. The document contains two separate sheets. Dataset C corresponds to 6,101 prePAAR-containing sequences that were identified through an iterative search of the UniprotKB using Tse6_NT_. Dataset D corresponds to 1,166 sequences collected following filtering of dataset C (see methods for details).

**Table 3. Summary of the number of genomes and effector sequences used in our informatics analyses (provided as Table_S3_methods_D01.xlsx file)**. This document contains three separate sheets. The “UniprotKB-effectors” sheet shows the quantity of initial prePAAR-containing sequences that were identified in our search and the number of sequences that were used following filtering and removal of redundancy (plotted in the cladogram in Figure S1A). The numbers in bold indicate the number of sequences in Table 1. The “8 genera -genomes” sheet corresponds to the number of genomes from the 8 genera (*Burkholderia*, *Escherichia*, *Enterobacter*, *Pseudomonas, Salmonella*, *Serratia, Shigella* and *Yersinia*) that contained one prePAAR-containing sequence and the number that remained following filtering and removal of redundancy. The “8-genera – effectors” sheet corresponds to initial and final numbers of prePAAR-containing sequences that were identified in the 8 genera listed above. The final number of sequences in this sheet were used to construct the cladogram in Figure 1E. The numbers in bold indicate the numbers of sequences in the datasets in Table 2.

## Supplementary figures

**Figure 1—figure supplement 1.**
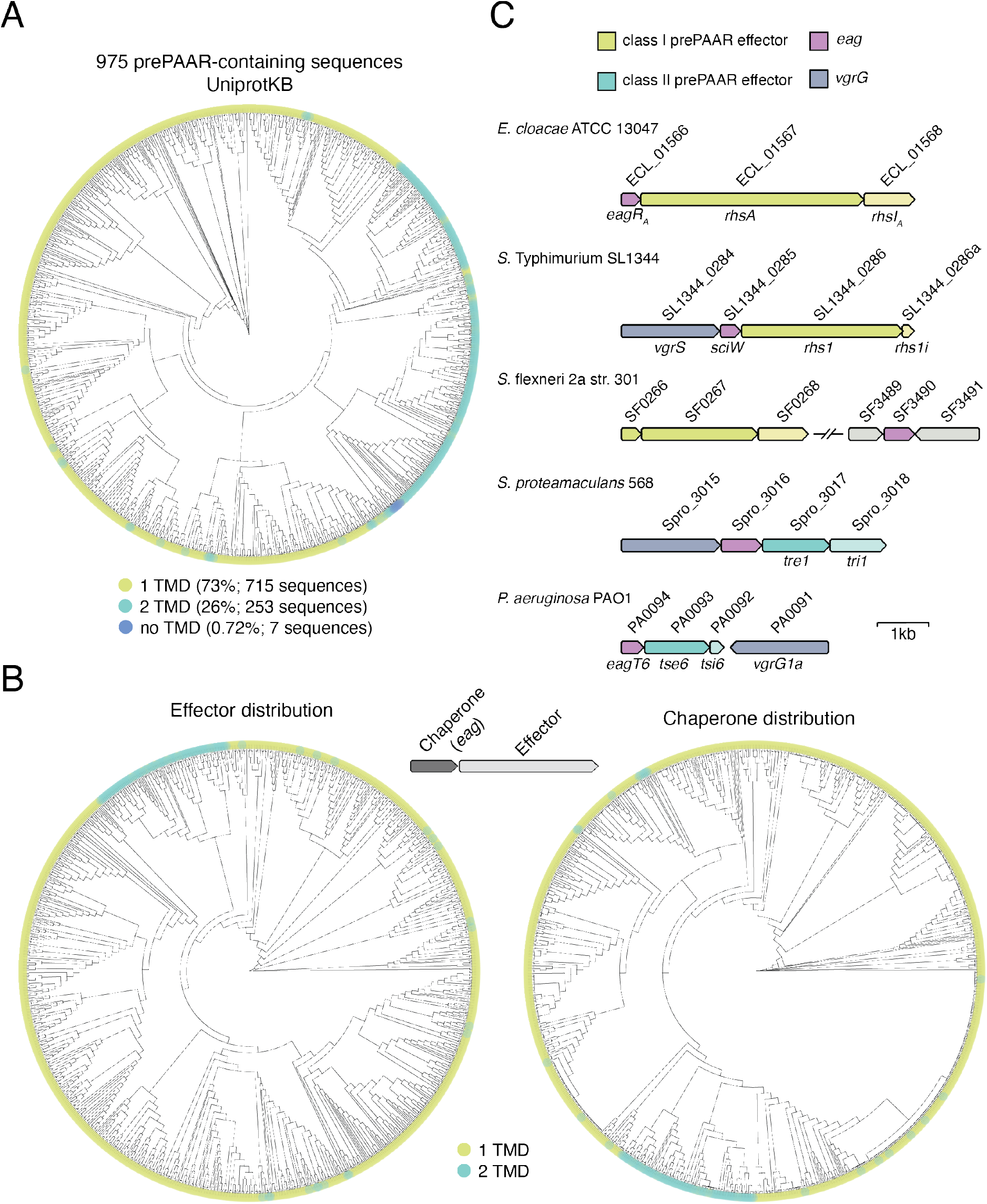
prePAAR effectors contain a fixed number of transmembrane domains. A) Phylogenetic distribution of 975 prePAAR-containing proteins identified in the UniProtKB database using the N-terminus of Tse6 (Tse6_NT_) as a search query (see methods). The TM helix predictors TMHMM and Phobius (Krogh et al., 2001, Kall et al., 2007) were used to quantify the number of TMDs in each protein (green, 1 TMD; blue, 2 TMDs). B) Similar analysis as Figure 1E, except that only prePAAR-containing effectors with an adjacent *eag* gene are depicted (left). The adjacently encoded *eag* chaperone sequences for each prePAAR effector were then used to build a second tree to depicting their distribution and association with an effector class (right). The *eag* chaperones were labelled with their neighbouring effector’s TMD prediction. All branch length represents evolutionary distance. C) Genomic arrangement of the five chaperone-effector pairs used for the co-purification experiment shown in Figure 1G. Shading was used to differentiate effector (dark) from potential immunity (light) genes. Locus tags and previously established names for each open reading frame are indicated above and below the gene diagram, respectively. Scale bar indicates 1 kilobase pair.

**Figure 2—figure supplement 1.**
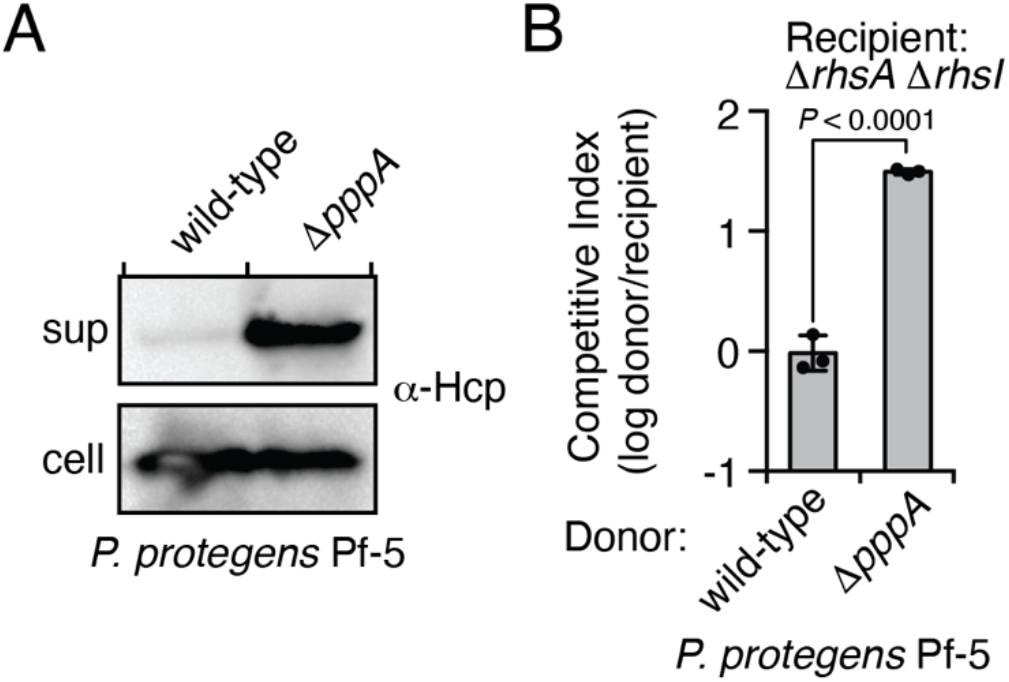
The type VI secretion system of *P. protegens* Pf-5 is repressed by the threonine phosphorylation pathway. A) Western blot of supernatant (sup) and cell fractions of the indicated *P. protegens* Pf-5 strains grown to OD 0.8. An Hcp (PFL_6089)-specific antibody was used to assess T6SS activity. B) Intraspecific growth competition assay of the indicated donor *P. protegens* strains against a recipient susceptible to intoxication by the class I prePAAR effector RhsA. Data are mean ±s.d. for *n* = 3 biological replicates; *P* value shown is from a two-tailed, unpaired *t*-test.

**Figure 3—figure supplement 1.**
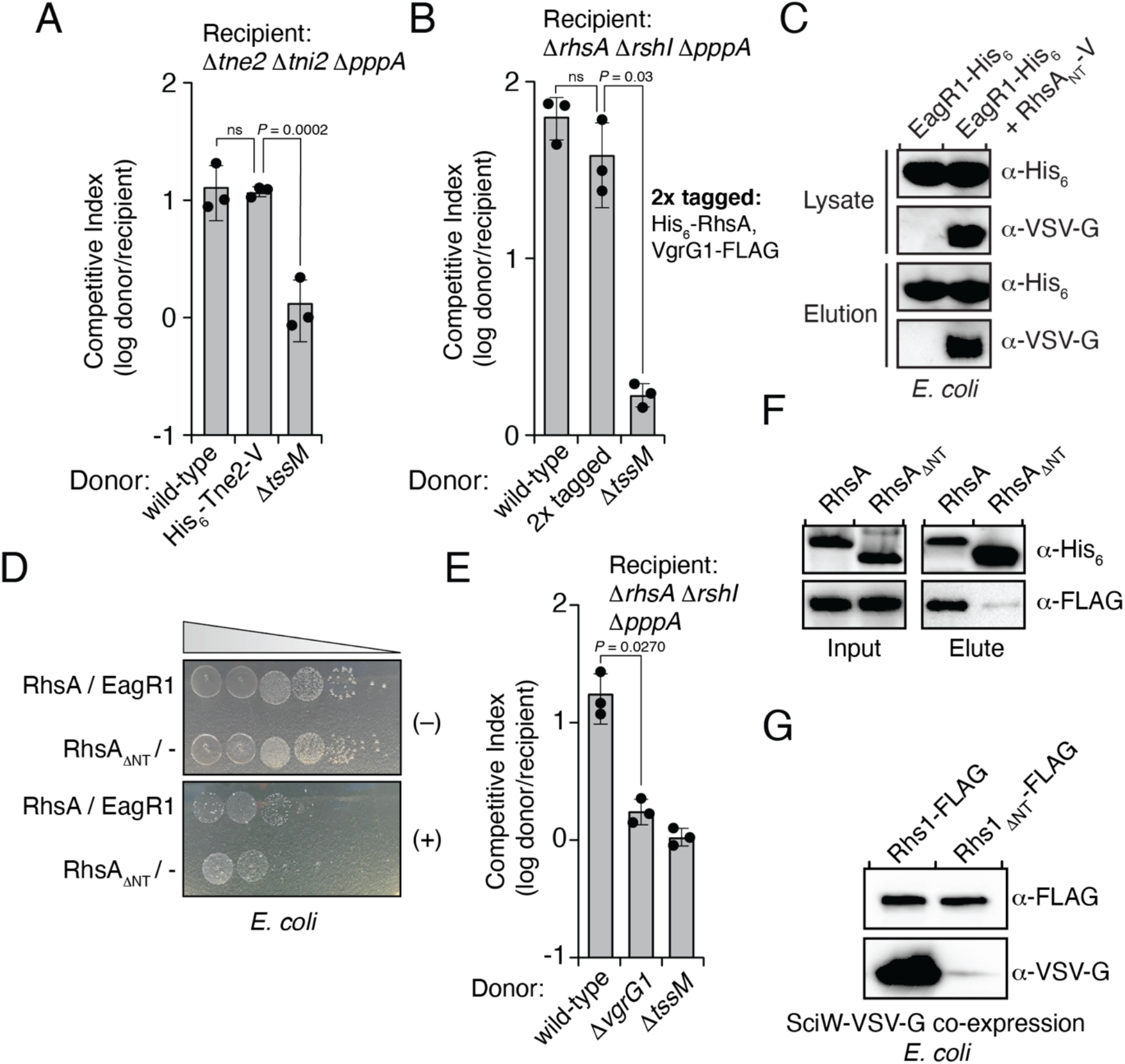
RhsA interacts with EagR1 and requires VgrG1 for delivery into target cells. A and B) Growth competition assays between the indicated *P. protegens* donor strains and either Tne2 (A) or RhsA (B) susceptible recipients. C) Western blot of lysate and pull-down elution fractions of His_6_-tagged EagR1 co-expressed with an empty vector or RhsA_NT_-VSV-G (residues 1-74) in *E. coli*. D) Growth of *E. coli* co-expressing inducible plasmids harboring RhsA and EagR1 or RhsA_ΔNT_ with an empty vector. Overnight cultures were plated on media containing (+) or lacking (−) inducers and were imaged after 24h of growth. E) Competition assay of the indicated *P. protegens* donor strains against a recipient susceptible to RhsA. F) Affinity pull-down of His_6_-tagged RhsA or RhsA_ΔNT_ co-expressed with VgrG-FLAG and EagR1-VSV-G in *E. coli*. Samples were analysed by western blot using the indicated antibodies. G) Western blot of affinity pull-down elution fractions of His_6_-tagged Rhs1 or Rhs1_ΔNT_ co-expressed with VSV-G tagged SciW. A-B, E) Data are mean ±s.d. for *n* = 3 biological replicates; *P* values shown are from a two-tailed, unpaired *t*-test. C-D, F-G) Data are representative of two independent experiments.

**Figure 4—figure supplement 1.**
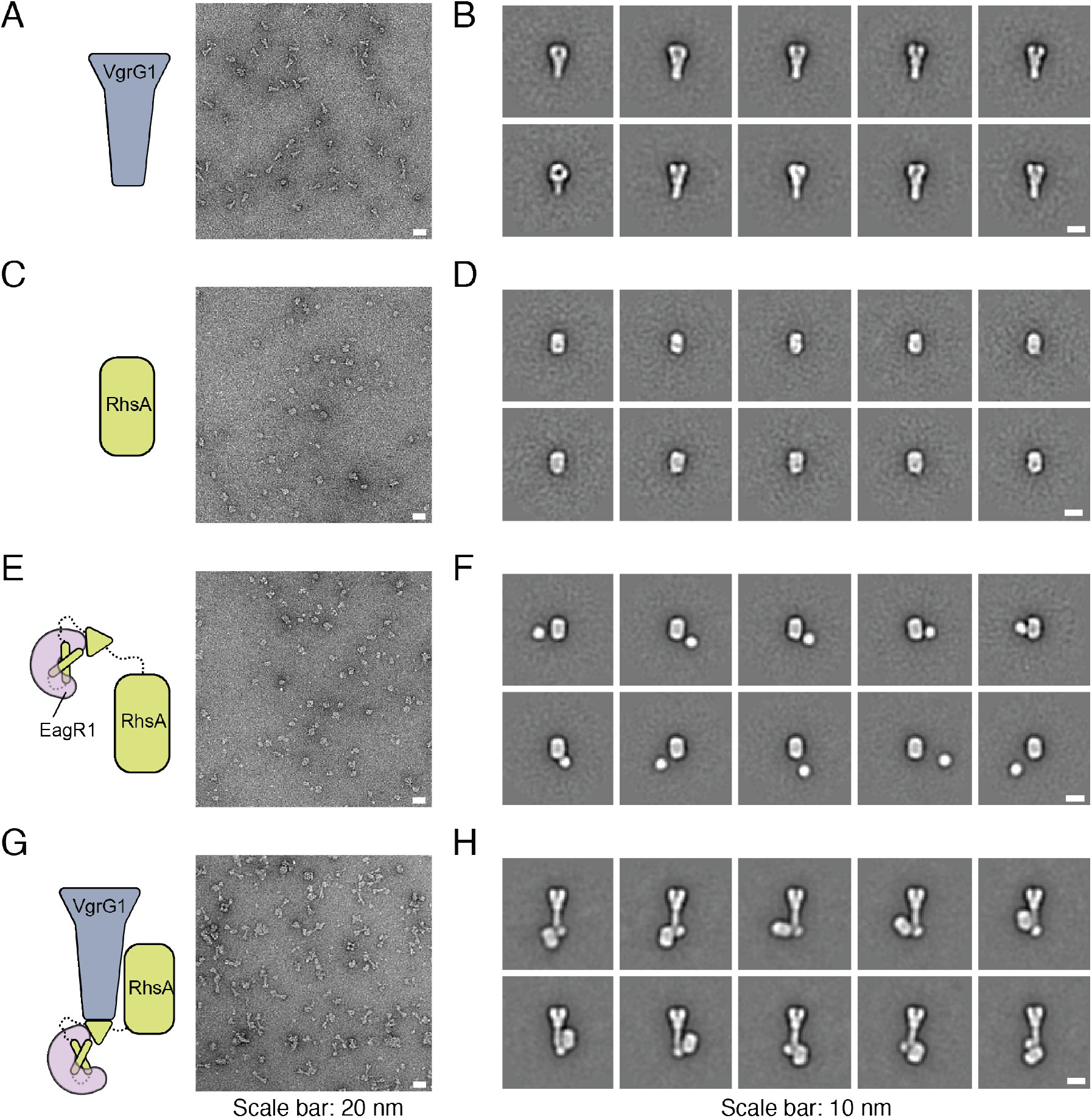
RhsA, EagR1 and VgrG1 form a ternary complex in vitro. Unprocessed micrographs (A, C, E, G) and representative 2-D class averages (B, D, F, H) of negatively stained VgrG1 (A, B), RhsA_ΔNT_ (C, D), EagR1-RhsA complex (E, F) and EagR1-RhsA-VgrG1 complex (G, H). Scale bar represents 20 nm for unprocessed micrographs and 10 nm for class averages.

**Figure 5—figure supplement 1.**
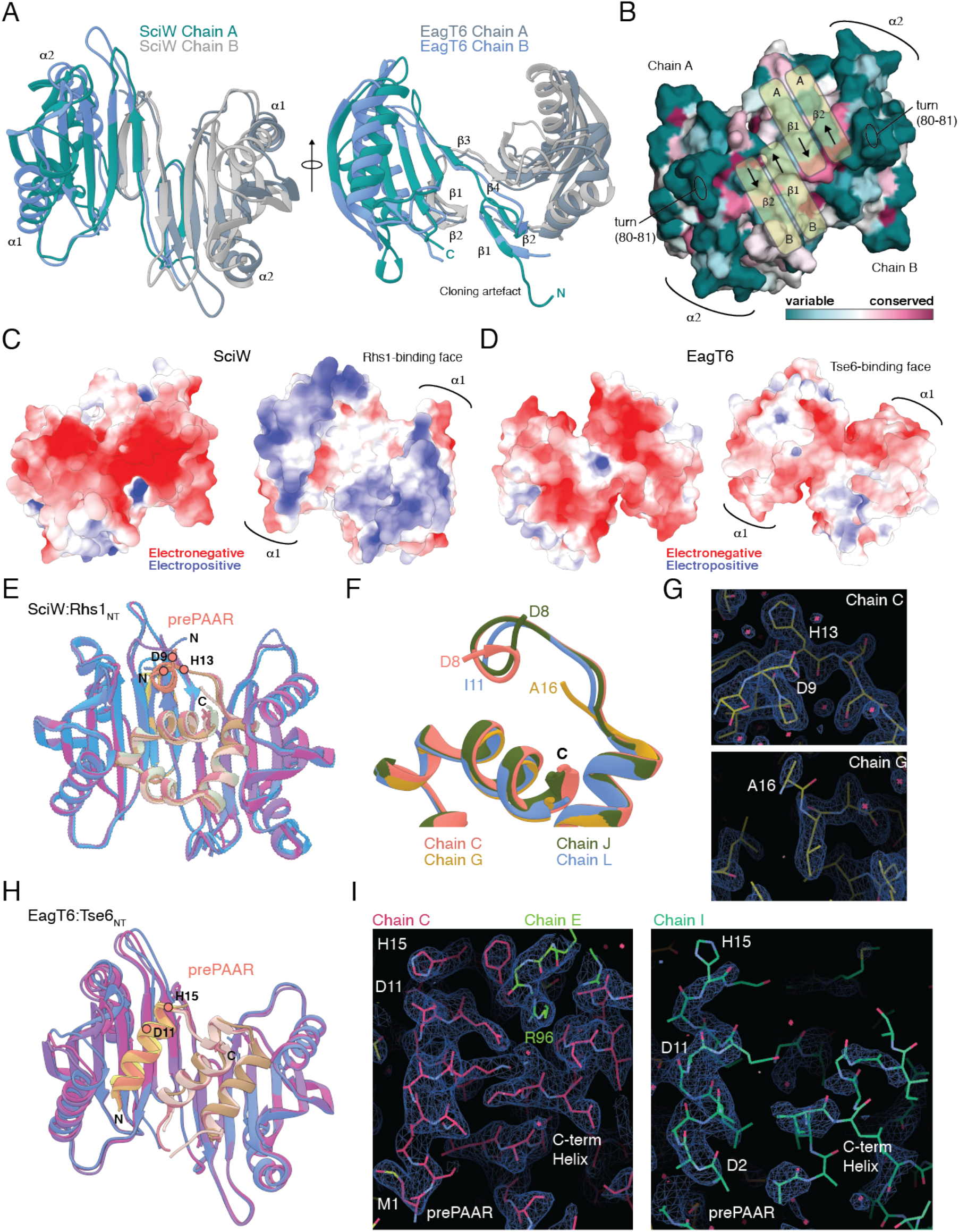
Structural comparison of Eag chaperones and effector complexes. A) Structural comparison of apo-SciW and apo-EagT6. Two views are shown related by an ~90° rotation. Each chaperone is colored by chain as in Figure 4. B) Conserved surface residues as determined by the Consurf server. The view is a 180° rotation of panel A from Figure 4. The domain-swap created by the beta-strands from chain A and chain B are labeled and shown with yellow bar overlays. C) Electrostatic surface potential of apo-SciW. The back (left, same surface as panel B) and Rhs1 binding surfaces (right) are shown. D) Electrostatic surface potential of apo-SciW. The convex (left, same surface as panel B) and concave (Tse6 binding) surfaces (right) are shown. E) Structural overlay of the four SciW-Rhs1_NT_ complexes in the asymmetric unit of the crystal structure. The modeled prePAAR and C-terminus of Rhs1 are indicated and colored by chain. F) View of the Rhs1 prePAAR region of each complex in the crystal structure. The N-terminal residue for each chain is listed. G) Electron density maps of SciW-Rhs1_NT_ Chain C and Chain G contoured at 1.4 rmsd (0.6816e/Å^3^). H) Structural overlay of the three EagT6-Tse6_NT_ complexes in the asymmetric unit of the crystal structure. The modeled prePAAR and C-terminus of Tse6 are indicated and colored by chain. I) Electron density maps of EagT6-Tse6_NT_ Chain C and Chain I contoured at 1.2 rmsd (0.0344e/Å^3^). The prePAAR and modelled C-terminal helix of the TMD region are labeled. A crystal packing artefact from Chain E including residue R96 that locks the prePAAR-TMD into place is shown. Electrostatic surface potentials were calculated by the adaptive-Poisson-Boltzmann server. Potentials are colored from −5 to 5 kT/e at pH 7.0. Images were created using UCSF Chimera, Coot, and Pymol.

**Figure 6—figure supplement 1.**
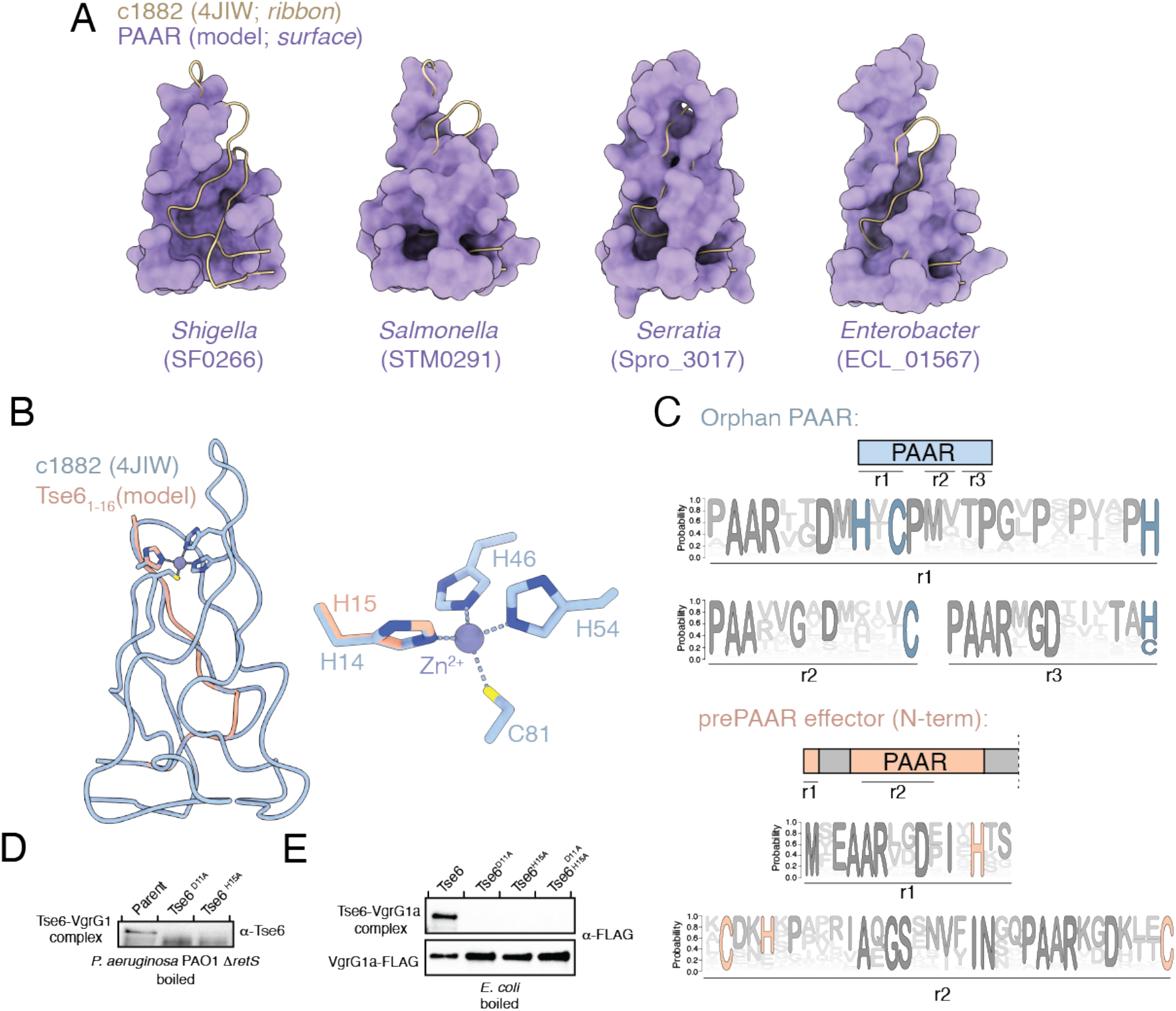
The PAAR domain of prePAAR effectors lacks a critical N-terminal segment. A) Surface representation of structural models of the PAAR domain from each of the indicated prePAAR effector proteins (purple) overlaid with a ribbon representation of the c1882 PAAR protein crystal structure (beige). Structural alignments were performed using ChimeraX. B) Structural overlay of the prePAAR segment (peach) from the artificially fused Tse6_prePAAR+PAAR_ sequence in Figure 6D with the entire c1882 PAAR protein (blue). The zoom shows the Zn^2+^-coordinating residues of c1882 and the overlap of H15 from Tse6’s prePAAR with H14 of c1882. C) Sequence logos developed from multiple sequence alignments of 564 orphan PAAR sequences and the N-terminus of 1,765 prePAAR-containing effectors. Sequence logos were developed for different regions (r1, r2, r3) in each construct that were contained for Zn^2+^-coordinating residues histidine and cysteine. D) Same samples from Figure 6A, except samples were boiled before being subject to electrophoresis. E) Same samples from 6G, except samples were boiled before being subject to electrophoresis.

## Notes

### Competing Interest Statement

The authors have declared no competing interest.

## References

Adamian, L. & Liang, J. 2002. Interhelical hydrogen bonds and spatial motifs in membrane proteins: polar clamps and serine zippers. Proteins, 47, 209–18.

Adams, P. D., Afonine, P. V., Bunkoczi, G., Chen, V. B., Davis, I. W., Echols, N., Headd, J. J., Hung, L. W., Kapral, G. J., Grosse-Kunstleve, R. W., Mccoy, A. J., Moriarty, N. W., Oeffner, R., Read, R. J., Richardson, D. C., Richardson, J. S., Terwilliger, T. C. & Zwart, P. H. 2010. PHENIX: a comprehensive Python-based system for macromolecular structure solution. Acta Crystallogr D Biol Crystallogr, 66, 213–21.

Ahmad, S., Wang, B., Walker, M. D., Tran, H. R., Stogios, P. J., Savchenko, A., Grant, R. A., Mcarthur, A. G., Laub, M. T. & Whitney, J. C. 2019. An interbacterial toxin inhibits target cell growth by synthesizing (p)ppApp. Nature.

Alcoforado Diniz, J. & Coulthurst, S. J. 2015. Intraspecies Competition in Serratia marcescens Is Mediated by Type VI-Secreted Rhs Effectors and a Conserved Effector-Associated Accessory Protein. J Bacteriol, 197, 2350–60.

Ashkenazy, H., Abadi, S., Martz, E., Chay, O., Mayrose, I., Pupko, T. & Ben-TAL, N. 2016. ConSurf 2016: an improved methodology to estimate and visualize evolutionary conservation in macromolecules. Nucleic Acids Res, 44, W344–50.

Basler, M., Ho, B. T. & Mekalanos, J. J. 2013. Tit-for-tat: type VI secretion system counterattack during bacterial cell-cell interactions. Cell, 152, 884–94.

Baynham, P. J., Ramsey, D. M., Gvozdyev, B. V., Cordonnier, E. M. & Wozniak, D. J. 2006. The Pseudomonas aeruginosa ribbon-helix-helix DNA-binding protein AlgZ (AmrZ) controls twitching motility and biogenesis of type IV pili. J Bacteriol, 188, 132–40.

Bondage, D. D., Lin, J. S., Ma, L. S., Kuo, C. H. & Lai, E. M. 2016. VgrG C terminus confers the type VI effector transport specificity and is required for binding with PAAR and adaptor-effector complex. Proc Natl Acad Sci U S A, 113, E3931–40.

Brooks, T. M., Unterweger, D., Bachmann, V., Kostiuk, B. & Pukatzki, S. 2013. Lytic activity of the Vibrio cholerae type VI secretion toxin VgrG-3 is inhibited by the antitoxin TsaB. J Biol Chem, 288, 7618–25.

Burkinshaw, B. J., Liang, X., Wong, M., Le, A. N. H., Lam, L. & Dong, T. G. 2018. A type VI secretion system effector delivery mechanism dependent on PAAR and a chaperone-co-chaperone complex. Nat Microbiol, 3, 632–640.

Busby, J. N., Panjikar, S., Landsberg, M. J., Hurst, M. R. & Lott, J. S. 2013. The BC component of ABC toxins is an RHS-repeat-containing protein encapsulation device. Nature, 501, 547–50.

Capella-Gutierrez, S., Silla-Martinez, J. M. & Gabaldon, T. 2009. trimAl: a tool for automated alignment trimming in large-scale phylogenetic analyses. Bioinformatics, 25, 1972–3.

Cardona, S. T. & Valvano, M. A. 2005. An expression vector containing a rhamnose-inducible promoter provides tightly regulated gene expression in Burkholderia cenocepacia. Plasmid, 54, 219–28.

Chen, V. B., Arendall, W. B., 3rd, Headd, J. J., Keedy, D. A., Immormino, R. M., Kapral, G. J., Murray, L. W., Richardson, J. S. & Richardson, D. C. 2010. MolProbity: all-atom structure validation for macromolecular crystallography. Acta Crystallogr D Biol Crystallogr, 66, 12–21.

Cianfanelli, F. R., Alcoforado Diniz, J., Guo, M., De Cesare, V., Trost, M. & Coulthurst, S. J. 2016a. VgrG and PAAR Proteins Define Distinct Versions of a Functional Type VI Secretion System. PLoS Pathog, 12, e1005735.

Cianfanelli, F. R., Monlezun, L. & Coulthurst, S. J. 2016b. Aim, Load, Fire: The Type VI Secretion System, a Bacterial Nanoweapon. Trends Microbiol, 24, 51–62.

Curran, A. R. & Engelman, D. M. 2003. Sequence motifs, polar interactions and conformational changes in helical membrane proteins. Curr Opin Struct Biol, 13, 412–7.

Dawson, J. P., Melnyk, R. A., Deber, C. M. & Engelman, D. M. 2003. Sequence context strongly modulates association of polar residues in transmembrane helices. J Mol Biol, 331, 255–62.

Dawson, J. P., Weinger, J. S. & Engelman, D. M. 2002. Motifs of serine and threonine can drive association of transmembrane helices. J Mol Biol, 316, 799–805.

Emsley, P., Lohkamp, B., Scott, W. G. & Cowtan, K. 2010. Features and development of Coot. Acta Crystallogr D Biol Crystallogr, 66, 486–501.

Flaugnatti, N., Le, T. T., Canaan, S., Aschtgen, M. S., Nguyen, V. S., Blangy, S., Kellenberger, C., Roussel, A., Cambillau, C., Cascales, E. & Journet, L. 2016. A phospholipase A1 antibacterial Type VI secretion effector interacts directly with the C-terminal domain of the VgrG spike protein for delivery. Mol Microbiol, 99, 1099–118.

Galan, J. E. & Waksman, G. 2018. Protein-Injection Machines in Bacteria. Cell, 172, 1306–1318.

Goodman, A. L., Kulasekara, B., Rietsch, A., Boyd, D., Smith, R. S. & Lory, S. 2004. A signaling network reciprocally regulates genes associated with acute infection and chronic persistence in Pseudomonas aeruginosa. Dev Cell, 7, 745–54.

Hmelo, L. R., Borlee, B. R., Almblad, H., Love, M. E., Randall, T. E., Tseng, B. S., Lin, C., Irie, Y., Storek, K. M., Yang, J. J., Siehnel, R. J., Howell, P. L., Singh, P. K., Tolker-Nielsen, T., Parsek, M. R., Schweizer, H. P. & Harrison, J. J. 2015. Precision-engineering the Pseudomonas aeruginosa genome with two-step allelic exchange. Nat Protoc, 10, 1820–41.

Hood, R. D., Singh, P., Hsu, F., Guvener, T., Carl, M. A., Trinidad, R. R., Silverman, J. M., Ohlson, B. B., Hicks, K. G., Plemel, R. L., Li, M.,Schwarz, S., Wang, W. Y., Merz, A. J., Goodlett, D. R. & Mougous, J. D. 2010. A type VI secretion system of Pseudomonas aeruginosa targets a toxin to bacteria. Cell Host Microbe, 7, 25–37.

Hyatt, D., Chen, G. L., Locascio, P. F., Land, M. L., Larimer, F. W. & Hauser, L. J. 2010. Prodigal: prokaryotic gene recognition and translation initiation site identification. BMC Bioinformatics, 11, 119.

Johnson, L. S., Eddy, S. R. & Portugaly, E. 2010. Hidden Markov model speed heuristic and iterative HMM search procedure. BMC Bioinformatics, 11, 431.

Jones, P., Binns, D., Chang, H. Y., Fraser, M., Li, W., Mcanulla, C., Mcwilliam, H., Maslen, J., Mitchell, A., Nuka, G., Pesseat, S., Quinn, A. F., Sangrador-Vegas, A., Scheremetjew, M., Yong, S. Y., Lopez, R. & Hunter, S. 2014. InterProScan 5: genome-scale protein function classification. Bioinformatics, 30, 1236–40.

Kabsch, W. 2010. Xds. Acta Crystallogr D Biol Crystallogr, 66, 125–32.

Kall, L., Krogh, A. & Sonnhammer, E. L. 2007. Advantages of combined transmembrane topology and signal peptide prediction--the Phobius web server. Nucleic Acids Res, 35, W429–32.

Katoh, K. & Standley, D. M. 2013. MAFFT multiple sequence alignment software version 7: improvements in performance and usability. Mol Biol Evol, 30, 772–80.

Kelley, L. A., Mezulis, S., Yates, C. M., Wass, M. N. & Sternberg, M. J. 2015. The Phyre2 web portal for protein modeling, prediction and analysis. Nat Protoc, 10, 845–58.

Krampen, L., Malmsheimer, S., Grin, I., Trunk, T., Luhrmann, A., De Gier, J. W. & Wagner, S. 2018. Revealing the mechanisms of membrane protein export by virulence-associated bacterial secretion systems. Nat Commun, 9, 3467.

Krogh, A., Larsson, B., Von Heijne, G. & Sonnhammer, E. L. 2001. Predicting transmembrane protein topology with a hidden Markov model: application to complete genomes. J Mol Biol, 305, 567–80.

Lacourse, K. D., Peterson, S. B., Kulasekara, H. D., Radey, M. C., Kim, J. & Mougous, J. D. 2018. Conditional toxicity and synergy drive diversity among antibacterial effectors. Nat Microbiol, 3, 440–446.

Li, W. & Godzik, A. 2006. Cd-hit: a fast program for clustering and comparing large sets of protein or nucleotide sequences. Bioinformatics, 22, 1658–9.

Liang, X., Moore, R., Wilton, M., Wong, M. J., Lam, L. & Dong, T. G. 2015. Identification of divergent type VI secretion effectors using a conserved chaperone domain. Proc Natl Acad Sci U S A, 112, 9106–11.

Mariano, G., Trunk, K., Williams, D. J., Monlezun, L., Strahl, H., Pitt, S. J. & Coulthurst, S. J. 2019. A family of Type VI secretion system effector proteins that form ion-selective pores. Nat Commun, 10, 5484.

Mok, B. Y., De MORAES, M. H., Zeng, J., Bosch, D. E., Kotrys, A. V., Raguram, A., Hsu, F., Radey, M. C., Peterson, S. B., Mootha, V. K., Mougous, J. D. & Liu, D. R. 2020. A bacterial cytidine deaminase toxin enables CRISPR-free mitochondrial base editing. Nature.

Moriya, T., Saur, M., Stabrin, M., Merino, F., Voicu, H., Huang, Z., Penczek, P. A., Raunser, S. & Gatsogiannis, C. 2017. High-resolution Single Particle Analysis from Electron Cryo-microscopy Images Using SPHIRE. J Vis Exp.

Mougous, J. D., Cuff, M. E., Raunser, S., Shen, A., Zhou, M., Gifford, C. A., Goodman, A. L., Joachimiak, G., Ordonez, C. L., Lory, S., Walz, T., Joachimiak, A. & Mekalanos, J. J. 2006. A virulence locus of Pseudomonas aeruginosa encodes a protein secretion apparatus. Science, 312, 1526–30.

Mougous, J. D., Gifford, C. A., Ramsdell, T. L. & Mekalanos, J. J. 2007. Threonine phosphorylation post-translationally regulates protein secretion in Pseudomonas aeruginosa. Nat Cell Biol, 9, 797–803.

Murshudov, G. N., Vagin, A. A. & Dodson, E. J. 1997. Refinement of macromolecular structures by the maximum-likelihood method. Acta Crystallogr D Biol Crystallogr, 53, 240–55.

Nguyen, V. S., Jobichen, C., Tan, K. W., Tan, Y. W., Chan, S. L., Ramesh, K., Yuan, Y., Hong, Y., Seetharaman, J., Leung, K. Y., Sivaraman, J. & Mok, Y. K. 2015. Structure of AcrH-AopB Chaperone-Translocator Complex Reveals a Role for Membrane Hairpins in Type III Secretion System Translocon Assembly. Structure, 23, 2022–31.

Paulsen, I. T., Press, C. M., Ravel, J., Kobayashi, D. Y., Myers, G. S. A., Mavrodi, D. V., Deboy, R. T., Seshadri, R., Ren, Q., Madupu, R., Dodson, R. J., Durkin, A. S., Brinkac, L. M., Daugherty, S. C., Sullivan, S. A., Rosovitz, M. J., Gwinn, M. L., Zhou, L., Schneider, D. J., Cartinhour, S. W., Nelson, W. C., Weidman, J., Watkins, K., Tran, K., Khouri, H., Pierson, E. A., Pierson, L. S., Thomashow, L. S. & Loper, J. E. 2005. Complete genome sequence of the plant commensal Pseudomonas fluorescens Pf-5. Nature Biotechnology, 23, 873–878.

Pei, T. T., Li, H., Liang, X., Wang, Z. H., Liu, G., Wu, L. L., Kim, H., Xie, Z., Yu, M., Lin, S., Xu, P. & Dong, T. G. 2020. Intramolecular chaperone-mediated secretion of an Rhs effector toxin by a type VI secretion system. Nat Commun, 11, 1865.

Pettersen, E. F., Goddard, T. D., Huang, C. C., Couch, G. S., Greenblatt, D. M., Meng, E. C. & Ferrin, T. E. 2004. UCSF Chimera—a visualization system for exploratory research and analysis. J Comput Chem, 25, 1605–12.

Pissaridou, P., Allsopp, L. P., Wettstadt, S., Howard, S. A., Mavridou, D. A. I. & Filloux, A. 2018. The Pseudomonas aeruginosa T6SS-VgrG1b spike is topped by a PAAR protein eliciting DNA damage to bacterial competitors. Proc Natl Acad Sci U S A, 115, 12519–12524.

Price, M. N., Dehal, P. S. & Arkin, A. P. 2010. FastTree 2--approximately maximum-likelihood trees for large alignments. PLoS One, 5, e9490.

Quentin, D., Ahmad, S., Shanthamoorthy, P., Mougous, J. D., Whitney, J. C. & Raunser, S. 2018. Mechanism of loading and translocation of type VI secretion system effector Tse6. Nat Microbiol, 3, 1142–1152.

Renault, M. G., Zamarreno Beas, J., Douzi, B., Chabalier, M., Zoued, A., Brunet, Y. R., Cambillau, C., Journet, L. & Cascales, E. 2018. The gp27-like Hub of VgrG Serves as Adaptor to Promote Hcp Tube Assembly. J Mol Biol, 430, 3143–3156.

Rietsch, A., Vallet-GELY, I., Dove, S. L. & Mekalanos, J. J. 2005. ExsE, a secreted regulator of type III secretion genes in Pseudomonas aeruginosa. Proc Natl Acad Sci U S A, 102, 8006–11.

Russell, A. B., Hood, R. D., Bui, N. K., Leroux, M., Vollmer, W. & Mougous, J. D. 2011. Type VI secretion delivers bacteriolytic effectors to target cells. Nature, 475, 343–7.

Samsonov, V. V., Samsonov, V. V. & Sineoky, S. P. 2002. DcrA and dcrB Escherichia coli genes can control DNA injection by phages specific for BtuB and FhuA receptors. Res Microbiol, 153, 639–46.

Shneider, M. M., Buth, S. A., Ho, B. T., Basler, M., Mekalanos, J. J. & Leiman, P. G. 2013. PAAR-repeat proteins sharpen and diversify the type VI secretion system spike. Nature, 500, 350–353.

Silverman, J. M., Agnello, D. M., Zheng, H., Andrews, B. T., Li, M., Catalano, C. E., Gonen, T. & Mougous, J. D. 2013. Haemolysin coregulated protein is an exported receptor and chaperone of type VI secretion substrates. Mol Cell, 51, 584–93.

Spinola-Amilibia, M., Davo-Siguero, I., Ruiz, F. M., Santillana, E., Medrano, F. J. & Romero, A. 2016. The structure of VgrG1 from Pseudomonas aeruginosa, the needle tip of the bacterial type VI secretion system. Acta Crystallogr D Struct Biol, 72, 22–33.

Tang, G., Peng, L., Baldwin, P. R., Mann, D. S., Jiang, W., Rees, I. & Ludtke, S. J. 2007. EMAN2: an extensible image processing suite for electron microscopy. J Struct Biol, 157, 38–46.

Tang, J. Y., Bullen, N. P., Ahmad, S. & Whitney, J. C. 2018. Diverse NADase effector families mediate interbacterial antagonism via the type VI secretion system. J Biol Chem, 293, 1504–1514.

Tareen, A. & Kinney, J. B. 2020. Logomaker: beautiful sequence logos in Python. Bioinformatics, 36, 2272–2274.

Ting, S. Y., Bosch, D. E., Mangiameli, S. M., Radey, M. C., Huang, S., Park, Y. J., Kelly, K. A., Filip, S. K., Goo, Y. A., Eng, J. K., Allaire, M., Veesler, D., Wiggins, P. A., Peterson, S. B. & Mougous, J. D. 2018. Bifunctional Immunity Proteins Protect Bacteria against FtsZ-Targeting ADP-Ribosylating Toxins. Cell, 175, 1380–1392 e14.

Ting, S. Y., Martinez-Garcia, E., Huang, S., Bertolli, S. K., Kelly, K. A., Cutler, K. J., Su, E. D., Zhi, H., Tang, Q., Radey, M. C., Raffatellu, M., Peterson, S. B., De LORENZO, V. & Mougous, J. D. 2020. Targeted Depletion of Bacteria from Mixed Populations by Programmable Adhesion with Antagonistic Competitor Cells. Cell Host Microbe.

Unterweger, D., Kostiuk, B. & Pukatzki, S. 2017. Adaptor Proteins of Type VI Secretion System Effectors. Trends Microbiol, 25, 8–10.

Vance, R. E., Rietsch, A. & Mekalanos, J. J. 2005. Role of the type III secreted exoenzymes S, T, and Y in systemic spread of Pseudomonas aeruginosa PAO1 in vivo. Infect Immun, 73, 1706–13.

Wagner, T., Merino, F., Stabrin, M., Moriya, T., Antoni, C., Apelbaum, A., Hagel, P., Sitsel, O., Raisch, T., Prumbaum, D., Quentin, D., Roderer, D., Tacke, S., Siebolds, B., Schubert, E., Shaikh, T. R., Lill, P., Gatsogiannis, C. & Raunser, S. 2019. SPHIRE-crYOLO is a fast and accurate fully automated particle picker for cryo-EM. Commun Biol, 2, 218.

Whitney, J. C., Beck, C. M., Goo, Y. A., Russell, A. B., Harding, B. N., De Leon, J. A., Cunningham, D. A., Tran, B. Q., Low, D. A., Goodlett, D. R., Hayes, C. S. & Mougous, J. D. 2014. Genetically distinct pathways guide effector export through the type VI secretion system. Mol Microbiol, 92, 529–42.

Whitney, J. C., Quentin, D., Sawai, S., Leroux, M., Harding, B. N., Ledvina, H. E., Tran, B. Q., Robinson, H., Goo, Y. A., Goodlett, D. R., Raunser, S. & Mougous, J. D. 2015. An interbacterial NAD(P)(+) glycohydrolase toxin requires elongation factor Tu for delivery to target cells. Cell, 163, 607–19.

Winn, M. D., Ballard, C. C., Cowtan, K. D., Dodson, E. J., Emsley, P., Evans, P. R., Keegan, R. M., Krissinel, E. B., Leslie, A. G., Mccoy, A., Mcnicholas, S. J., Murshudov, G. N., Pannu, N. S., Potterton, E. A., Powell, H. R., Read, R. J., Vagin, A. & Wilson, K. S. 2011. Overview of the CCP4 suite and current developments. Acta Crystallogr D Biol Crystallogr, 67, 235–42.

Winn, M. D., Isupov, M. N. & Murshudov, G. N. 2001. Use of TLS parameters to model anisotropic displacements in macromolecular refinement. Acta Crystallogr D Biol Crystallogr, 57, 122–33.

Wood, T. E., Howard, S. A., Wettstadt, S. & Filloux, A. 2019. PAAR proteins act as the 'sorting hat' of the type VI secretion system. Microbiology, 165, 1203–1218.

Yang, Z., Fang, J., Chittuluru, J., Asturias, F. J. & Penczek, P. A. 2012. Iterative stable alignment and clustering of 2D transmission electron microscope images. Structure, 20, 237–47.

Yu, G. 2020. Using ggtree to Visualize Data on Tree-Like Structures. Curr Protoc Bioinformatics, 69, e96.

Zhang, D., Iyer, L. M. & Aravind, L. 2011. A novel immunity system for bacterial nucleic acid degrading toxins and its recruitment in various eukaryotic and DNA viral systems. Nucleic Acids Res, 39, 4532–52.

